# Bacterial Pathogen Infection Triggers Magic Spot Nucleotide Signaling in *Arabidopsis thaliana* Chloroplasts Through Specific RelA/SpoT Homologs

**DOI:** 10.1101/2023.04.26.538375

**Authors:** Danye Qiu, Esther Riemer, Thomas M. Haas, Isabel Prucker, Shinji Masuda, Yan L. Wang, Georg Felix, Gabriel Schaaf, Henning J. Jessen

## Abstract

Magic spot nucleotides (p)ppGpp are important signaling molecules in bacteria and plants. In the latter, RSH enzymes are responsible for (p)ppGpp turnover. Profiling of (p)ppGpp is more difficult in plants than in bacteria due to lower concentrations and more severe matrix effects. Here, we report that capillary electrophoresis mass spectrometry (CE-MS) can be deployed to study (p)ppGpp abundance and identity in *Arabidopsis thaliana*. This goal is achieved by combining a titanium dioxide extraction protocol and pre-spiking with chemically synthesized stable isotope labeled internal reference compounds. The high sensitivity and separation efficiency of CE-MS enables monitoring of changes in (p)ppGpp levels in *A. thaliana* upon infection with the pathogen *Pseudomonas syringae* pv. *tomato (PstDC3000)*. We observed a significant increase of ppGpp post infection that is also stimulated by the flagellin peptide flg22 only. This increase depends on functional flg22 receptor FLS2 and its interacting kinase BAK1 indicating that pathogen-associated molecular pattern (PAMP) receptor-mediated signaling controls ppGpp levels. Transcript analyses showed an upregulation of *RSH2* upon flg22 treatment and both *RSH2* and *RSH3* after *PstDC3000* infection. *A. thaliana* mutants deficient in RSH2 and RSH3 activity display no ppGpp accumulation upon infection and flg22 treatment, supporting involvement of these synthases in PAMP-triggered innate immunity responses to pathogens within the chloroplast.

## Introduction

The total global biomass (estimated to ca. 550 gigatons of carbon (Gt C)) is dominated by plants (450 Gt C) followed by bacteria (70 Gt C).^1^ These kingdoms have developed various ways of interaction, exemplified by symbiotic plant microbiota interactions, such as nitrogen fixation in root nodules, or invasive competition as plant pathogens.^2, 3^ The endosymbiosis of cyanobacterial-like prokaryotes leading to the development of chloroplasts for photosynthesis is a key event marking the prelude to dominance of the plant kingdom on earth.^4^ Such a symbiotic fusion would likely require the interplay and harmonization of kingdom-specific signaling pathways

The bacterial stringent response (SR) to stress is governed by the magic spot nucleotides (p)ppGpp, densely phosphorylated guanosine nucleotides with a 5′-triphosphate or 5′-diphosphate moiety combined with a 3′-diphosphate group.^5^ Discovered more than 50 years ago,^6^ their diverse functions help bacteria cope with different stresses, most prominently amino acid starvation-mediated growth adjustment.^7^ While there are only very few reports on (p)ppGpp as a signaling entity in metazoa,^8, 9^ plants have retained the ability to generate (p)ppGpp within chloroplasts,^10–14^ with ppGpp as the by far most abundant representative (Magic Spot I, Figure 1).

**Figure 1.**
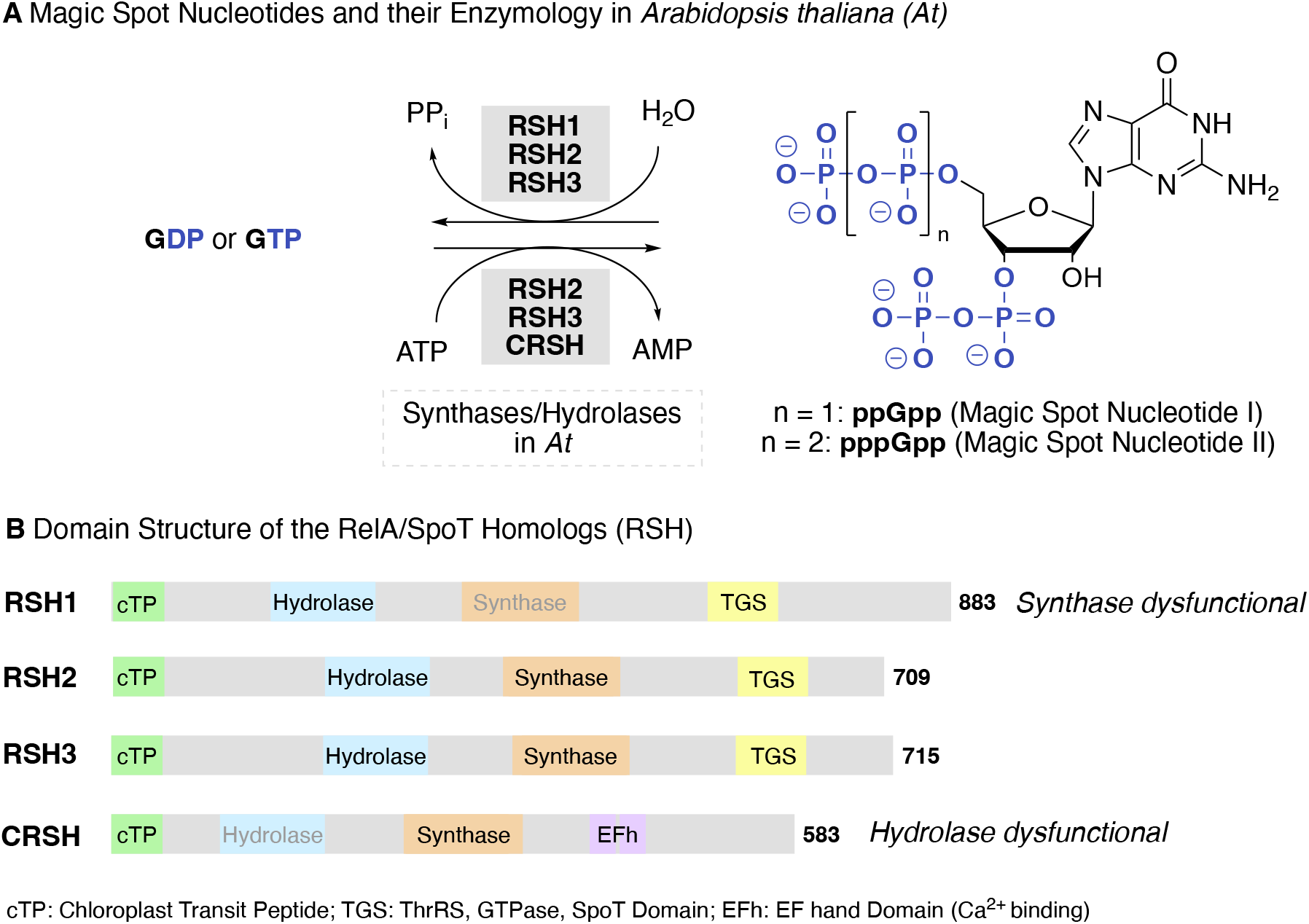
**A)** Structures of (p)ppGpp and different RelA/SpoT homologs (RSH enzymes) found in *A. thaliana* that are responsible for (p)ppGpp metabolism. **B)** Overview of the domain structures of the RSH enzymes in *A. thaliana*. The colored boxes represent domains and their location within proteins is roughly indicated. Domains written in grey indicate dysfunctionality of the domain. All homologs have an N-terminal chloroplast transit peptide (cTP). AtCRSH lacks the ThrRS, GTPase and SpoT (TGS) domain, but contains a Ca^2+^ binding EF hand motif (EFh).

There have been significant advances in the understanding of (p)ppGpp function and regulation in bacteria, in particular regarding protein interactomes.^15–19^ However, in plants, only comparatively little is known about (p)ppGpp signaling and the chloroplastic interactomes of ppGpp remain largely uncharacterized with few exceptions.^20^ Even so, it is now established that (p)ppGpp are important for plant adaptation to stress, regulation of chloroplast function, nitrogen starvation, and onset of immune responses,^14, 21–31^ which was summarized in a recent review.^32^

In the past decade, several members of the RelA-SpoT homolog (RSH) enzyme superfamily that antagonistically synthesize and/or hydrolyse (p)ppGpp were identified in plants and algae.^33, 34^ Four nuclear-encoded RSH enzymes are found in *Arabidopsis*: RSH1, RSH2, RSH3, and the Ca^2+^-dependent RSH (CRSH) (Figure 1B), all of which are suggested to localize to the chloroplasts via a chloroplast transit peptide.^12, 14, 22, 26^ They are multidomain proteins that contain synthase, hydrolase, and regulatory domains, however not all domains are functional. For example, RSH1 lacks an amino acid crucial for (p)ppGpp synthesis and therefore only has hydrolase activity.^21^ RSH2 and RSH3 share high amino-acid identity (80%) and are bifunctional enzymes with synthase and hydrolase activities, mostly responsible for ppGpp production during the day^22, 23, 26, 28^ and a CRSH counterbalancing hydrolase activity during night.^30^ CRSH functions exclusively as (p)ppGpp synthase.^30, 35^ The Ca^2+^-dependent homolog is suggested to mediate the Stringent Response (SR) by sensing and responding to calcium fluctuations via (p)ppGpp production, which might help plants to adapt to stresses, such as wounding and insect invasion.^22, 34^ CRSH responds to light-to-dark transition by transient (p)ppGpp synthesis.^30, 36^ Pathogen-associated molecular pattern (PAMP) receptor triggered immunity (PTI) *in planta* provokes comparable Ca^2+^ fluxes as are evoked by darkness, and thus CRSH might be activated during PTI.^37–39^ However, a control experiment treating *crsh* mutant plants with the PAMP activating peptide flagellin22 (flg22), a truncated 22 amino acid version of the full bacterial flagellin, still induced defense-related genes as in wildtype^30^ and therefore uncertainties remain regarding CRSH involvement in PTI.^32^ *RSH2 and RSH3* transcript levels are upregulated by plant pathogenic viruses^25^ as well as salicylic acid (SA).^13, 14, 40^ ppGpp levels are directly correlated with susceptibility to Turnip Mosaic Virus infection while there is an inverse correlation regarding salicylic acid (SA) responsive transcript levels of defense-related PR1 (PATHOGENESIS RELATED 1).^25^ Overall, it has become clear that ppGpp signaling is involved in the plant immune response but a full picture has not yet emerged.^10^ Here, we show that ppGpp accumulates under light in *A. thaliana* whole seedlings after treatment with the bacterial plant pathogen *Pseudomonas syringae* pv. *tomato (PstDC3000)*, potentially representing a defensive signaling response to bacterial infection. We demonstrate that ppGpp is produced by the plant – not the bacteria – through RSH2/3 in response to PAMPs, mediated in part by the flagellin receptor FLS2 and its interacting kinase BAK1. The required absolute ppGpp quantitation is achieved with a novel capillary electrophoresis mass spectrometry method (CE-MS) using synthetic heavy isotope labeled internal reference compounds and TiO_2_ enrichment to minimize matrix effects.

## Results and Discussion

The extraction and quantitation of Magic Spot Nucleotides poses significant challenges that have been mainly addressed in bacteria. Radioactive phosphate labeling and thin layer chromatography in the beginning^6^ has now been mostly superseded with liquid chromatography (LC) and ion chromatography (IC) mass-spectrometry-based approaches (MS), but also UV detection has been applied. ^14, 30, 36, 41, 42^ Double spike isotope dilution IC-MS has been introduced to correct for pppGpp decomposition during extraction in bacteria.^43^ The problems of decomposition during extraction and ion suppression due to matrix effects are aggravated if one switches from bacteria to the more complex plant matrices, even more so as the absolute concentration of ppGpp in plants is lower.^41^ In this regard, moving from UV to MS detection has led to a significant reduction of required plant material by ca. 200-fold. Today, plant tissue samples around 100 mg can be profiled routinely.^36^ The reliability of ppGpp quantitation by mass spectrometry in plants has been significantly improved by a recent publication from the Field laboratory.

They introduced enzymatically prepared stable heavy isotope internal (p)ppGpp reference compounds that can also correct for losses during extraction without separate control runs.^41^In an earlier study, we have shown that capillary electrophoresis (CE) is an alternative separation platform for densely phosphorylated nucleotides, such as ppGpp, relying on UV detection.^44^ Here, we demonstrate that CE coupled to electrospray ionization (ESI)-MS using a triple quadrupole system (QqQ) is a reliable alternative to LC-based methods. For a quantitative analysis, we introduce chemically synthesized stable heavy isotope (p)ppGpp reference compounds that are fully ^15^N labeled (M+5; Figure 2). The chemoenzymatic synthesis enabled ready access to heavy ppGpp on a 15 mg scale. While pppGpp was also synthesized, we did not detect significant amounts of it in any of the plant samples, in agreement with a study from the Field laboratory^41^ and therefore this internal reference was not further applied.

**Figure 2.**
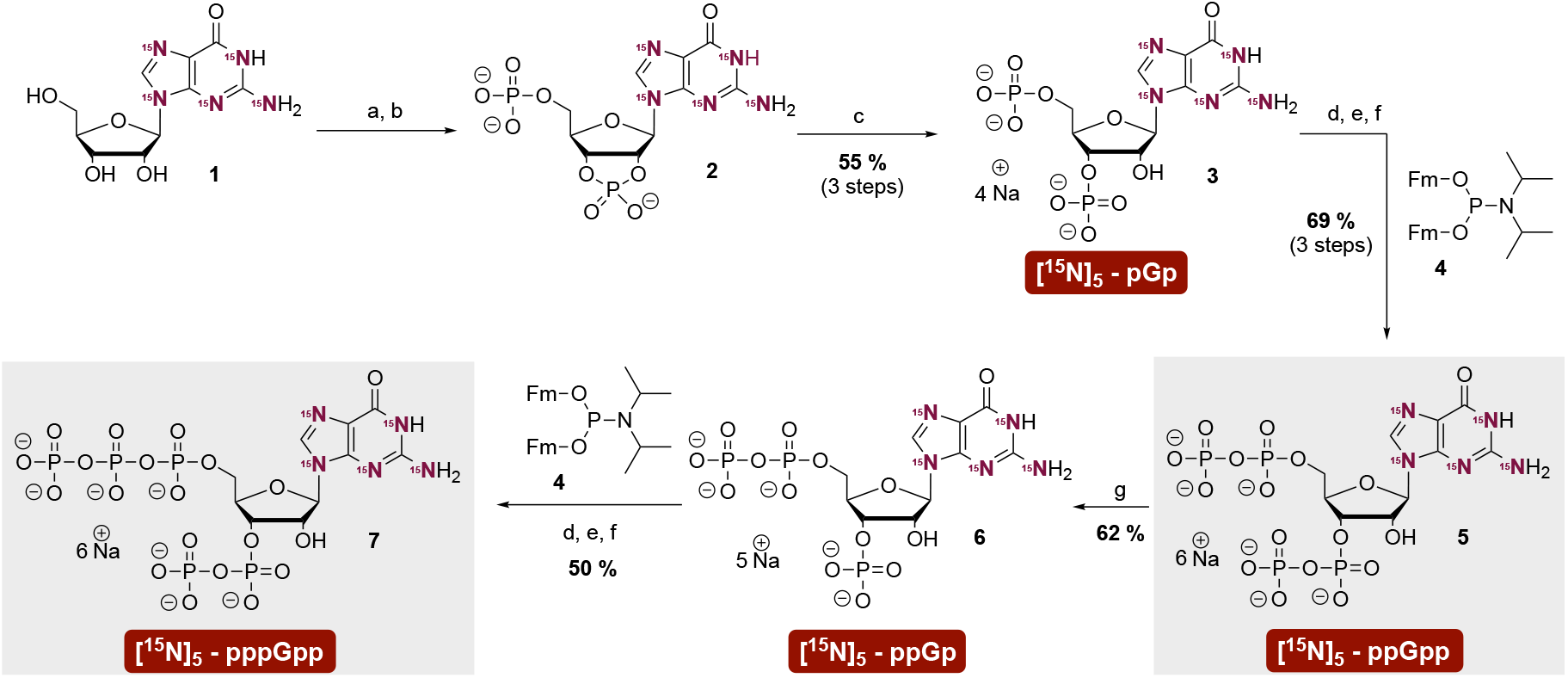
Synthesis of magic spot nucleotide [^15^N]_5_ - isotopologues. (a) P_2_Cl_4_O_3_ (20 eq.), 0 °C, 3 h. (b) NaHCO_3_ – buffer (1 M) (c) RNase T2, pH = 7.5, 37 °C, 12 h. (d) **4** (3 eq.), ETT (5 eq.), DMF, rt, 15 min. (e) *m*CPBA (3 eq.), -20 °C, 15 min. (f) DBU, rt, 30 min. (g) RNase T2, pH = 5.5, 37 °C, 12 h. Abbreviations: ETT 5-(ethylthio)-1H-tetrazole; *m*CPBA *meta-*chloroperbenzoic acid; DBU 1,8-diazabicyclo(5.4.0)undec-7-ene; rt room temperature; Fm fluorenylmethyl

### Chemical synthesis of heavy isotope labeled ppGpp

In brief, the synthesis commenced with commercially available ^15^N-labeled guanosine **1**. Treatment with pyrophosphorylchloride followed by partial hydrolysis gave cyclophosphate **2**. This intermediate was selectively ring-opened to pGp **3** by RNase T2.^45^ Both the 5′- and 3′-phosphates were homologated into diphosphates with bis-fluorenylmethyl P-amidite **4** ^46, 47^ giving access to ppGpp **5** on a multi-milligram scale. Treatment of labeled ppGpp **5** with RNase T2 at a pH of 5.5 led to 2′,3′ cyclophosphate formation followed by regioselective ring-opening to ppGp **6**, essentially corresponding to a selective 3′-pyrophosphatase reaction. ppGp **6** was then simultaneously homologated in the 5′- and 3′ positions with P-amidite **4** to give pppGpp **7**, again on a multi-milligram scale. Purification of the target molecules was achieved by strong anion exchange chromatography (SAX) with a sodium perchlorate eluent. Precipitation from cold acetone yielded the nucleotides as their sodium salts. Defined stock solutions of (p)ppGpp for CE-MS measurement were obtained using quantitative ^31^P-NMR or ^1^H-NMR spectroscopy. Detailed synthetic procedures and characterization data can be found in the supporting information (see supplementary information, sections 7–9).

### Extraction and CE-MS method development

Access to significant amounts of internal heavy isotope references enables spiking of nucleotides prior to extraction (pre-spiking). Pre-spiking enables correction for losses during extraction and precise quantitation irrespective of the matrix. During method development, we realized that the commonly used cold formic acid/SPE extraction for (p)ppGpp from plants was not suitable for CE measurements resulting in strong matrix effects and ion suppression. Therefore, we studied an alternative enrichment protocol that has been previously used to isolate inositol pyrophosphates from complex matrices for CE-MS measurements, including plant tissues. ^48–50^ In this approach, cell lysis is achieved with cold perchloric acid (PA), followed by pre-spiking with heavy internal references. Enrichment on TiO_2_ beads^51^ precedes elution with ammonium hydroxide solution. After evaporation and dissolution, the resulting extracts can then be analyzed by CE-MS. The entire sample preparation workflow is visualized in scheme 1, including the electropherogram of a separation of a reference nucleotide mixture using an optimized background electrolyte (ammonium acetate [35 mM, pH = 9.7]).

**Scheme 1.**
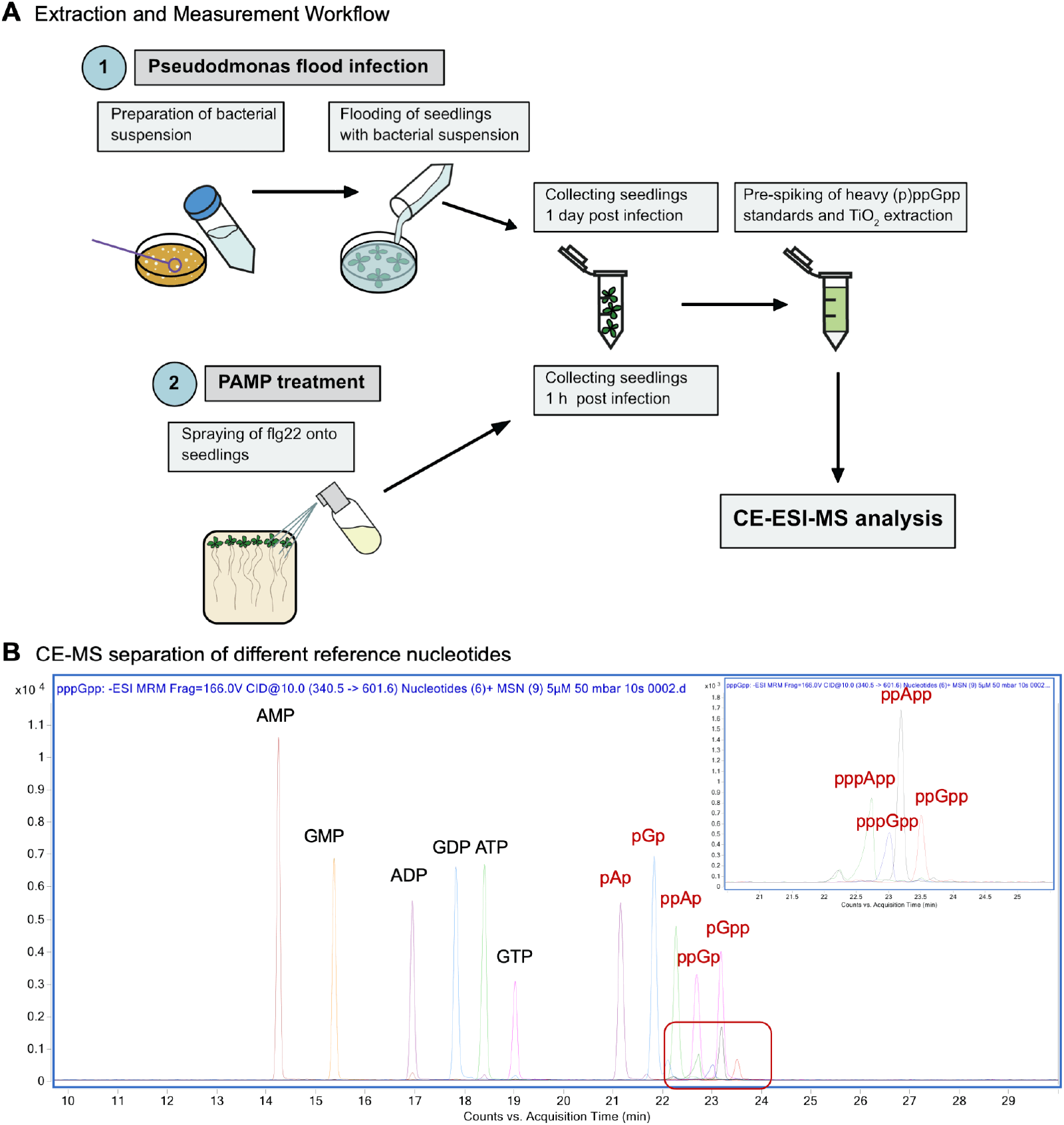
**A** Flow-chart of sample preparation for CE-ESI-MS analysis. Different treatments are discussed in the section on plant pathogen interactions below. **B** CE-MS measurement of a reference nucleotide mixture (5 μm stock solutions, injection volume 10 nL), Magic Spot Nucleotides highlighted in red, the red box is additionally magnified on the upper right.

We observed partial chemical decomposition of the analyte under the applied conditions: likely by transient formation of a 2′,3′-*cyclo*phosphate intermediate, the eluate from the TiO_2_ beads contained variable amounts of ppGp with a monophosphate in either the 2′ or 3′ position, indicative of non-selective chemical hydrolysis of the 2′,3′-*cyclo*phosphate. The putative products were validated by chemical synthesis and CE-MS analysis (see Supplementary Figures S1-4). The formation of these byproducts caused no problems, as pre-spiking was applied throughout this study ensuring correction for such losses. Additionally, the observed ppGp (2′) and ppGp (3′) are absent in all plant samples we analyzed and as a consequence, the absolute and relative abundance of these byproducts could be used for quantification and as an internal control. An example of an *A. thaliana* wildtype (Col-0) extract demonstrating these assignments is shown in the supporting information (see Supplementary Figure S1). Salient advantages of CE are high separation efficiencies in combination with nanoliter sample consumption and low operating costs. Importantly, the TiO_2_ extraction protocol will now enable to correlate ppGpp and InsP signaling in plants by parallel determinations, which will potentially uncover cross-talk between these pathways.^48, 50, 52-54^ Each analysis presented herein is based on a 20 nL injection volume of the analyte solution. Determination of the limit of detection (LOD) and limit of quantification (LOQ) were achieved via spiking of extracted plant material (150 mg fresh weight (FW)) with heavy internal standards and calculation of a signal to noise ratio above 3 (LOD) and 10 (LOQ). The LOD for ppGpp expressed as a concentration was 10 nM and the LOQ was 30 nM in this particular matrix (see Supplementary Figures S5 and S6). Calibration curves were linear and had a coefficient of determination >0.999 over the investigated range from 0.1 μM to 400 μM (see Supplementary Figure S7). In summary, we present a new extraction protocol streamlining quantitative analyses of ppGpp abundance from plant tissues by CE-MS using a triple quadrupole mass spectrometer.

### Plant pathogen interactions

With the CE-MS method available, we studied how plant infection by bacterial pathogens affects ppGpp levels. In this context, one must address the issue of ppGpp origin: plant or pathogen? Along these lines, we studied *Pseudomonas syringae* pv. *tomato (PstDC3000)* infected wildtype (Col-0) and mutant plants as well as specific PAMPs to stimulate PAMP triggered immunity (PTI) and avoid bacterial contaminations (see Scheme 1). Our results demonstrate that plants respond to bacterial pathogen infection in part via PTI by increasing ppGpp levels. In flagellin-treated plants, these increases are paralleled by upregulation of *RSH2* transcript levels. In plants infected with bacteria, increased ppGpp is paralleled by upregulation of *RSH2/RSH3* transcript levels.

We first determined the time frame in which the infection experiments could be run reliably as the pathogen will damage the plant over time. Inspection of the plants at various days post infection (dpi) showed serious damage after three days (Figure 3, A), the onset of disease after two days, whereas at 1 dpi, the plant still looked healthy. In parallel, we extracted infected *A. thaliana* seedlings (150 mg FW) 1, 2, or 3 dpi and analyzed ppGpp content by CE-MS (Figure 3, B) in the negative electrospray ionization mode (ESI^-^) using multiple reaction monitoring (MRM). We selected singly negatively charged ions (Figure 3, C) for quantitation. A significant increase of ppGpp was observed both at 1 and 2 dpi, whereas after three days also ppGpp in the mock control sample increased. We attribute this to the severe damage observed at 3 dpi. Since the signal increase was highly significant already 1 dpi (Figure 3, B and C), we continued our study with samples collected at 1 dpi.

**Figure 3.**
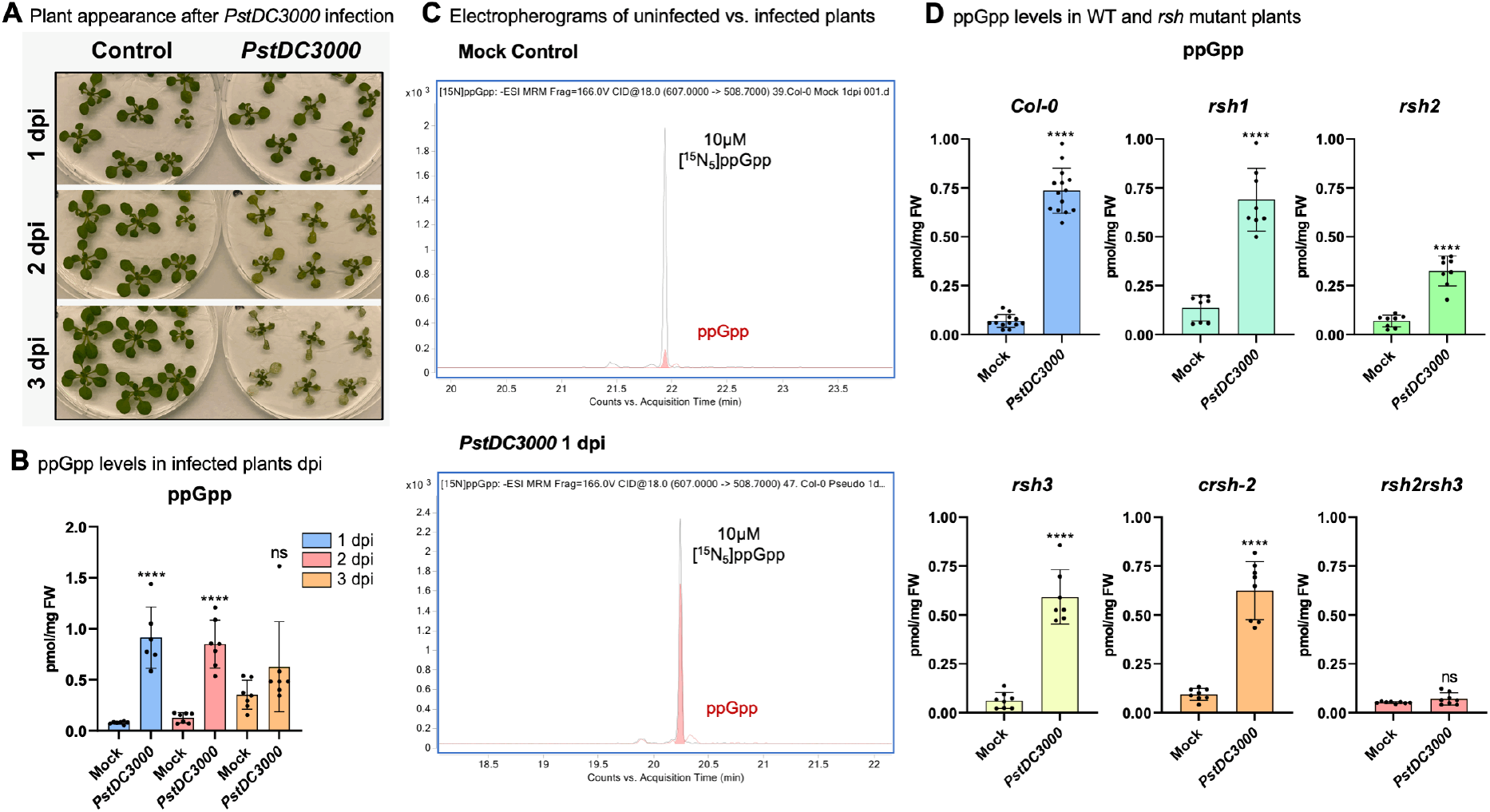
**A** Photographs of *Arabidopsis thaliana* seedlings taken after several days of infection with *Pseudomonas syringae pv. tomato (PstDC3000)*. **B** Quantitative analysis of ppGpp levels in *A. thaliana* seedlings 1, 2, or 3 days post infection (dpi) indicated as pmol per mg of fresh weight (FW). 150 mg of plant material was extracted under light. Data are presented as means ± SD (n = 7). **C** Examples of electropherograms obtained from CE-ESI-MS analysis with a QqQ analyzer in MRM mode. The recorded transition was 607.0 → 508.7 in negative ionization mode. Internal heavy references were spiked at 10 μM in either uninfected (mock) or infected samples (1 dpi). **D** Levels of ppGpp in wildtype (Col-0) versus single mutants (*rsh1, rsh2, rsh3, crsh-2*) or a double mutant (*rsh2 rsh3*) 1 day without infection (mock) or 1 dpi with *PstDC3000*. Data are presented as means ± SD (n = 7-14); ns = not significant; ****P < 0.0001, Student’s t-test.

Next, we analyzed ppGpp content in different *rsh* single (*rsh1, rsh2, rsh3, crsh-2*) and double (*rsh2 rsh3*) mutant plants. Seeds of T-DNA insertion lines *rsh2* (Sail_305_B12) and *rsh3* (Sail_99_G05), and *rsh1* (Sail_391_E11) were obtained from The European Arabidopsis Stock Centre (http://arabidopsis.info/) and Arabidopsis Biological Resource Center (https://abrc.osu.edu), respectively. The, *crsh-2* (CRISPR-modified *crsh*-null mutant) and the *rsh2 rsh3* double mutant (SAIL CS81411 x GABI 129D0) were described previously.^26,30^ Importantly, under non-stressed conditions, these plants grow healthy (see Supplementary Figure S8). Analysis of ppGpp levels enabled us to validate our method against established LC-MS protocols.^30, 36, 41^ Our results are in a similar range compared to data previously reported for wildtype and mutant plants without infection, thus validating our TiO_2_ extraction CE-MS workflow (Scheme 1 and Figure 3, B, D). For example, Bartoli et al.^41^ reported 28.7 ± 2.2 pmol g^−1^ for Col-0 *A. thaliana* whereas Ono et al.^30^ found 172.9 ± 15.6 pmol g^−1^. We recorded an intermediate value of 69.8 ± 33.2 pmol g^−1^ ppGpp. For *crsh-2*, Ono et al.^30^ found 186.8 ± 5.6 pmol g^−1^ of ppGpp, relatively close to the value of 94.0 ± 29.7 measured in this study. Figure 3 D summarizes the results of the analysis. Wildtype (Col-0) mock-treated samples have an average ppGpp content of 0.07 ± 0.03 pmol mg^−1^ fresh weight that increases by ca. 11-fold one day after infection. Mock-treated *rsh1* plants displayed higher ppGpp levels compared to Col-0 in line with a hydrolase-only activity of RSH1. *Pseudomonas*-infected *rsh1* plants showed an approx.5-fold increase in ppGpp levels. The *rsh2* mutant displayed wildtype levels of ppGpp and also increased ppGpp content upon infection, albeit to a lower degree as found for Col-0. Similar findings were made for the *rsh3* and the *crsh-2* single mutant, respectively, but the reduction of ppGpp production was less pronounced in these lines as found in *rsh2* mutant plants. Critically, the *rsh2 rsh3* double mutant does not accumulate ppGpp upon infection, showing that RSH2 and RSH3 have partially redundant functions and can compensate for loss of either one’s activity. We conclude that the increase in ppGpp upon infection is a result of RSH2 and/or RSH3 abundance/activity.

In plants, extracellular signal perception and transmission is regulated by leucine-rich repeat receptor kinases.^55^ The flagellin receptor (FLS2) recognizes flagellin, but is also sensitive to a truncated version of it, the elicitor-active epitope peptide flagellin 22 (flg22, Figure 4, A).^56^ Upon binding, FLS2 heterodimerizes with the receptor-like kinase BAK1, that phosphorylates downstream targets. The flg22 variant flg^Atum^ (Figure 4, A) does not induce downstream signaling via FLS2/BAK1.^57^ To investigate, whether ppGpp signaling in chloroplasts is mediated by PAMP recognition, we studied, if the flg22 peptide elicits ppGpp increases upon 1 hour stimulation. Mock and flg^Atum^ treatment served as negative controls. We conducted these studies in wildtype (Col-0) and in different mutant backgrounds as described above. Additionally, we tested *fls2 bak1-4* mutant plants^56^ that host a set of fully functional RSH enzymes (Figure 4, B). Relative to the mock control, treatment with flg^Atum^ did not result in increased ppGpp levels in any of the samples studied. However, 10 μM flg22 treatment for one hour led to a substantial increase (ca. 2-fold) of ppGpp in Col-0, and also in *rsh1, rsh3*, and *crsh-2*. The *rsh2* single mutant and *rsh2 rsh3* double mutant lost their ability to respond to flg22 with increases of ppGpp. Moreover, the *fls2 bak1-4* mutant also was not responsive to flg22 treatment anymore regarding increases in ppGpp. These data suggest that flg22 elicits increases in chloroplastic ppGpp levels by signaling mediated via PAMP receptor FLS2 and BAK1, eventually resulting in increased RSH2 activity and/or abundance. Consequently, we profiled transcript levels of the relevant *RSH* genes by qPCR in response to flg22 and also *PstDC3000* infection. The results are summarized in Figure 4 C.

**Figure 4.**
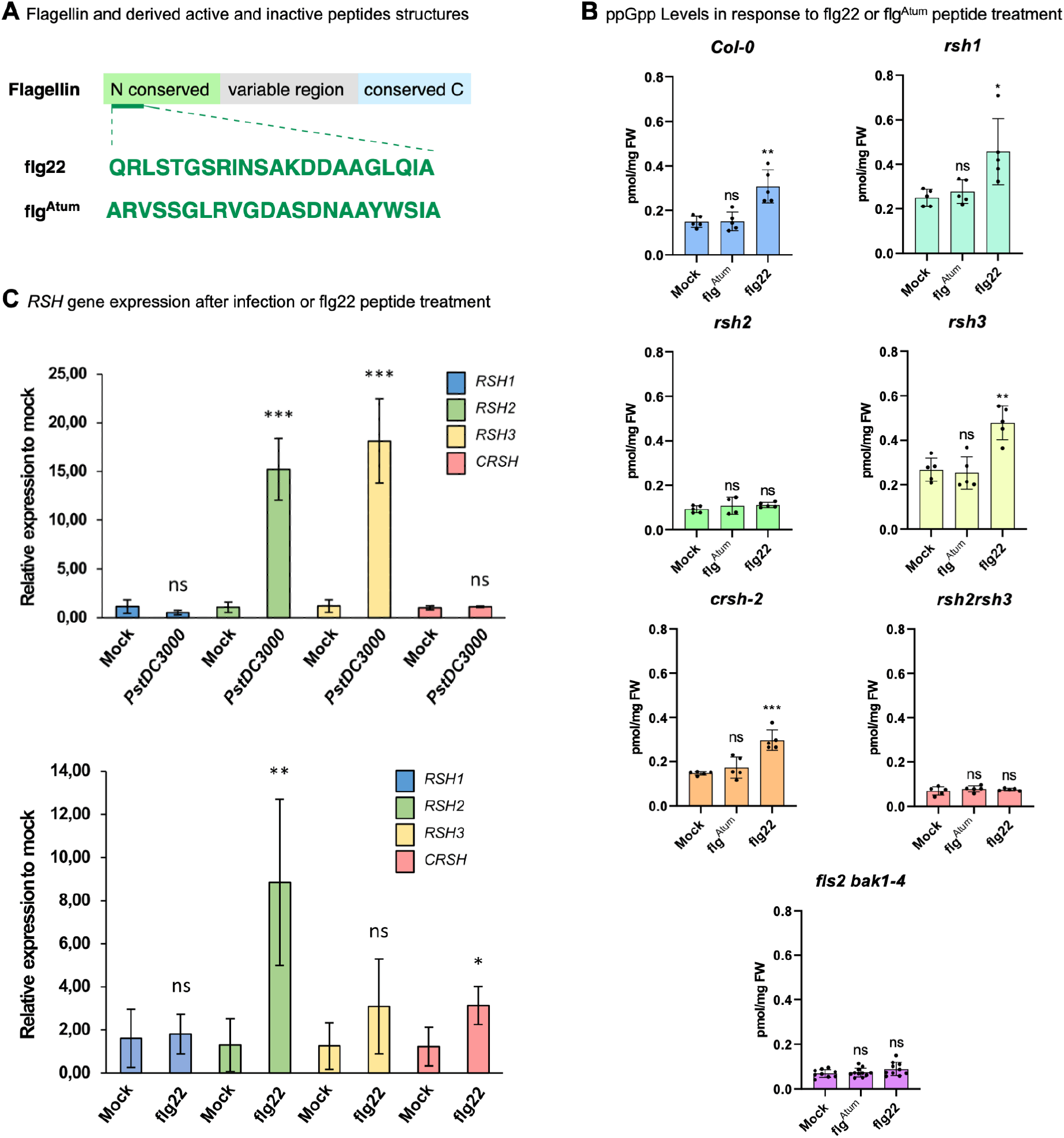
**A** Domain structure of flagellin and sequences of derived peptides (flg22 and flg^Atum^). **B** Quantitative analysis of ppGpp levels in *A. thaliana* seedlings 1 hour after treatment with peptides (10 μM) or mock control indicated as pmol mg^-1^ of fresh weight (FW). Data are presented as means ± SD (n ≥ 5). **C** qPCR analysis of gene expression 1 day after infection with *PstDC3000* (upper panel) or treatment for 1h with 10 μM flg22 (lower panel). Data are presented as means ± SD (n = 4); ns = not significant; *P < 0.05, **P < 0.01, ***P < 0.001, Student’s t-test.

Previously, it had been shown that plants react to infection by upregulation of stringent response genes. In particular, in tobacco infected with *Erwinia carotovora*^*13*^ or *Pectobacterium atrosepticum* an increase in *RSH2* transcript levels was recorded, whereas this was not the case in potato.^58^ Our results demonstrate that flg22 treatment triggers a ca. 8-fold upregulation of *RSH2* transcript compared to mock control. Transcript levels of other RSHs were not significantly affected with this treatment, except for *CRSH*, which showed a moderate upregulation (approx. 3-fold). This is in line with the failure of *rsh2* plants to increase ppGpp levels upon flg22 treatment. In comparison, both *RSH2* and *RSH3* transcripts were upregulated more than fifteenfold after infection with *PstDC3000*. This again is in line with the observation that only the double mutant *rsh2 rsh3* failed to increase ppGpp levels after infection with the pathogen *PstDC3000* (Figure 3, D). This in turn suggests that pathogen infection triggers a more diverse signaling response as compared to flg22 treatment alone, eventually leading to increased expression of both *RSH2* and *RSH3*.

## Conclusion

Plants have inherited the ability to synthesize ppGpp by bacterial and retained the magic spot nucleotide signaling pathway within their chloroplasts. In bacteria, these molecules govern the Stringent Response to stress. The pathways leading to increased ppGpp concentrations in plant chloroplasts and the signaling outcomes of such increases in plants remain largely uncharacterized. Several studies have highlighted the importance of ppGpp in plant/pathogen interactions, such as the regulation of ppGpp metabolism after viral infection^25^ or regulation of *RSH* gene expression after bacterial infection.^13, 58^ It was suggested that CRSH plays a role in such interactions, as Ca^2+^ signaling within the chloroplast occurs as a response to infection.^30^

In this study, we have developed a chemical synthesis of scalable multimilligram quantities of heavy isotope labeled internal (p)ppGpp reference compounds. These molecules were deployed to devise a highly sensitive CE-MS approach based on pre-spiking of the references using a new TiO_2_ extraction approach, enabling the absolute quantitation of the analytes from complex matrices. We applied this approach to study ppGpp produced by *A. thaliana* in response to bacterial *PstDC3000* infection. ppGpp levels increased ca. tenfold after one day post infection. While *rsh2 rsh3* double mutants showed no increase of ppGpp levels in response to infection, the single mutants still exhibited this response, indicating that loss of one RSH can be compensated, at least partially, by the other RSH enzyme. Notably, ppGpp increases were also elicited by treatment with the flg22 peptide that operates through activation of the receptor kinase FLS2. No induction was observed with flg^Atum^, the flg22 epitope variant of the crown gall disease causing *Agrobacterium tumefaciens*, that evades immunodetection by FLS2 of *A. thaliana*.^59^ Flg22-induced ppGpp increases, ca. two-fold, appeared weaker than after bacterial infection. However, the effect of flg22 was measured after 1 hour after incubation while the effect of bacterial infection was measured after 24 hours. In case of flg22 stimulation, single disruption of *RSH2* sufficed to block the response, whereas loss of *RSH3* did not suppress ppGpp increases. This suggests that PAMP signaling increases ppGpp levels through activation or increase of RSH2 enzymes. In contrast, a bacterial infection stimulates additional pathways that eventually activate or increase RSH2 and RSH3 enzymes simultaneously. Gene expression analyses by qPCR support this model, as both *RSH2* and *RSH3* transcripts are upregulated >15-fold after *Pseudomonas* infection, whereas flg22 treatment results in ca. 7-fold upregulation of the *RSH2* transcript level only.

Future studies will now have to address how precisely the involved signaling pathways, such as PTI, increase chloroplast ppGpp and to what avail the plant produces it. Here, interactome studies of ppGpp will likely provide new avenues for research. Additionally, such studies may help to understand how a ppGpp signal is relayed back to the nucleus, as suggested by increased *RSH* transcript levels.

## Acknowledgments

We thank Dr. Manfred Keller from MagRes of the University of Freiburg for a significant amount of time for NMR spectroscopy. This project has received funding from the European Research Council (ERC) under the European Union’s Horizon 2020 research and innovation program (grant agreement no. 864246, to H.J.J). This study was supported by the Deutsche Forschungsgemeinschaft (DFG) under Germany’s excellence strategy (CIBSS, EXC-2189, Project ID 390939984, to H.J.J. and EXC 2070 – 390732324, PhenoRob to G.S.) and a Grant-in-Aid for Scientific Research, KAKENHI (21H02075, to S.M.). We gratefully acknowledge financial support from the Studienstiftung des Deutschen Volkes, and the Brigitte Schlieben-Lange Programm.

