## Supplementary Information for "Bacterial Pathogen Infection Triggers Magic Spot Nucleotide Signaling in *Arabidopsis thaliana* Chloroplasts Through Specific RelA/SpoT Homologs"

**for**

#### 1. Bacterial strains and *Arabidopsis* T-DNA insertion lines used in this study

*Pseudomonas syringae* pv. *tomato* DC3000 (*Pst* DC3000). The strain was a gift from Florian Grundler (Molecular Phytomedicine, Institute of Crop Science and Resource Conservation, University of Bonn, Germany).

Col-0 wild type

*rsh1 Sail_391_E11*

*rsh2* Sail_305_B12

*rsh3* Sail_99_G05

*crsh-2* CRISPR-modified *crsh*-null mutant

*rsh2 rsh3* SAIL CS81411; GABI 129D0

*fls2 bak1-4* from Georg Felix (Center for Plant Molecular Biology Tübingen, Germany).

#### 2. ppGpp extraction from plants

Extraction of ppGpp from plant samples was carried out using titanium dioxide ($TiO_2$) enrichment method (rutil, Sigma Aldrich) as described previously (Mol. Plant 2021, 14, 1864). All steps were conducted at 4°C until elution. 5-10 mg $TiO_2$ beads were washed once in $H_2O$ and once in 1 M perchloric acid (PA). After washing, beads were supplemented with 300 pmol ppGpp and 30 pmol pppGpp standards. Plant material was homogenized by grinding the samples in liquid nitrogen. Homogenized plant material was mixed with PA and incubated for 10 min on ice. Samples were centrifuged twice at 18000 g to remove plant debris. Plant tissue was normalized to fresh weight by adjusting the volume of PA before adding the extracts to the $TiO_2$ beads. Samples were incubated at 4°C for 30 min, rotating.. After incubation, samples were centrifuged at 8000 g and the supernatants were discarded. $TiO_2$ beads were washed twice in PA before adding 200 µl of 10% $NH_4OH$ followed by incubation at room temperature for 5 min, rotating. Samples were spun and the supernatants were collected in fresh tubes. The step was repeated and the samples were vacuum evaporated using Speedvac for 2.5 h at 40°C.

#### 3. CE-ESI-QqQ

CE-ESI-QqQ analysis was performed on an Agilent 7100 CE System coupled with a triple quadrupole tandem mass spectrometry Agilent 6495c system, equipped with an Agilent Jet Stream (AJS) electrospray ionization (ESI) source. A sheath liquid coaxial interface from Agilent was employed, with an Agilent 1200 isocratic LC pump stably delivering the sheath-liquid (via a splitter set with a ratio of 1:100). The sheath liquid was composed of a water-isopropanol (1:1) mixture, which was introduced at a flow rate of 10 µL/min.

The experiments were conducted using a bare fused silica capillary with an internal diameter of 50 µm and a length of 100 cm. The background electrolyte (BGE) was 35 mM ammonium acetate, titrated to pH 9.7 with ammonia solution. A constant CE current of 22 µA was established by applying a voltage of +30 kV across the capillary. Samples were injected by applying a pressure of 100 mbar for 10 seconds (equivalent to 20 nL). The MS source parameters were set as follows: nebulizer pressure at 8 psi, gas temperature at 150 °C, sheath gas flow at 11 L/min and temperature at 175 °C, capillary voltage at -2000 V with nozzle voltage at 2000V. The negative high-pressure RF and low-pressure RF (Ion Funnel parameters) were set at 90V and 60V, respectively. The MassHunter Optimizer software identified the multiple reaction monitoring (MRM) transitions, and their corresponding mass spectrometer parameters are shown below.

| Compound | Precursor Ion | Product Ion | dwell | CE (V) | Cell Acc (V) | Polarity |
|---|---|---|---|---|---|---|
| [$^{15}N_5$]ppGpp | 607 | 508.7 | 80 | 18 | 3 | Negative |
| ppGpp | 602 | 503.7 | 80 | 18 | 3 | Negative |
| [$^{15}N_5$]pppGpp | 342.9 | 606.6 | 80 | 10 | 1 | Negative |
| pppGpp | 340.5 | 601.6 | 80 | 10 | 1 | Negative |
| [$^{15}N_5$]pppG | 527 | 429.1 | 80 | 21 | 3 | Negative |
| pppG | 522 | 424.1 | 80 | 21 | 3 | Negative |
| $IP_6$ | 328.9 | 78.8 | 50 | 46 | 3 | Negative |

##### 3.1 Analyte Assignment

Pre-spiking of $^{15}N$ labeled ppGpp enabled the assignment of decomposition products during $TiO_2$ enrichment. The following figure S4 shows how ppGpp partially decomposed to two isobaric analytes, which we assign as ppGp with a phosphate ester either in the 2´or 3´ position, designated as ppGp (2´); ppGp (3´). Given the same ratio of unlabeled and labeled analytes, we conclude that there is no natural ppGp in plants.

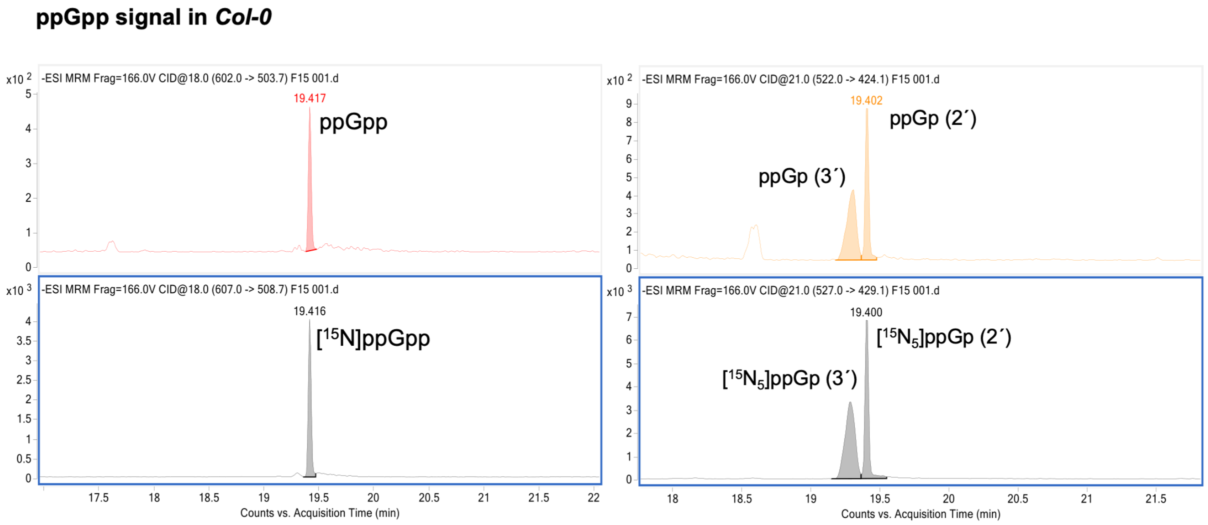


**Figure S1**: ppGpp and decomposition product signals after $TiO_2$ extraction of ppGpp from wild type plants (Col-0) with heavy isotope standard pre-spiking. Red trace: ppGpp of plant origin; orange trace: decomposition products derived from plant ppGpp during extraction. Grey traces: heavy ppGpp and ppGp as a result of concomitant heavy ppGpp degradation.

In order to unambiguously assign the decomposition products, we generated a mixture of heavy ppGp (2´and 3´) by basic treatment of $^{15}N$ labeled ppGpp, which leads to cyclophosphate formation to the required reference compounds. Figure S5 shows the clean conversion of the ppGpp reference to the desired mixture by removal of one phosphate unit ($P_i$). The coupled $^{31}P$ NMR also validates the regioselectivity.

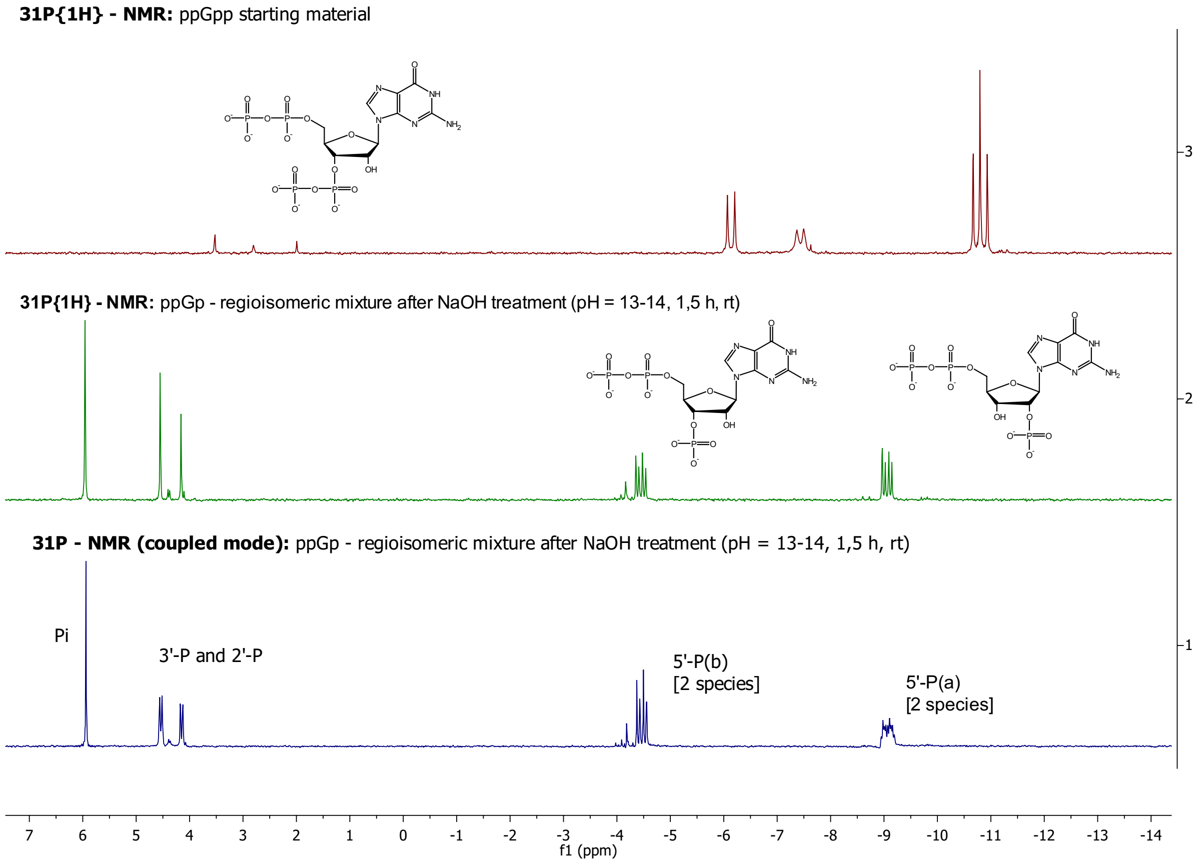


**Figure S2**: ppGpp hydrolysis leads to a regioisomeric mixture of ppGp (2´and 3´) as shown by $^{31}$P NMR spectroscopy.

The obtained mixture of ppGp (2´) and ppGp (3´) was analyzed by CE-MS, also including a synthetic sample of pGpp, which is another isobaric analyte. Indeed, the CE-MS protocol is capable to separate ppGp (2´) and ppGp (3´) and pGpp, as shown in figure S6. This underlines that our assignment of the $TiO_2$ decomposition products is correct.

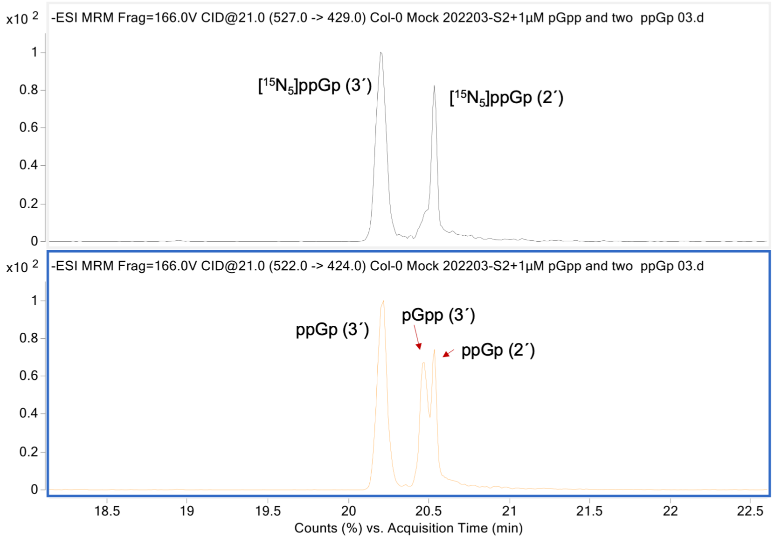


**Figure S3**: CE-MS analysis of heavy ppGpp regioisomeric mixture (top) and a regioisomeric mixture of ppGp (2´and 3´) with additionally spiked pGpp (3´).

Finally, we obtained high-resolution mass spectra of the analytes from plants after *Pseudomonas* infection (1dpi) using a Q-TOF system. These results further validate analyte identity.

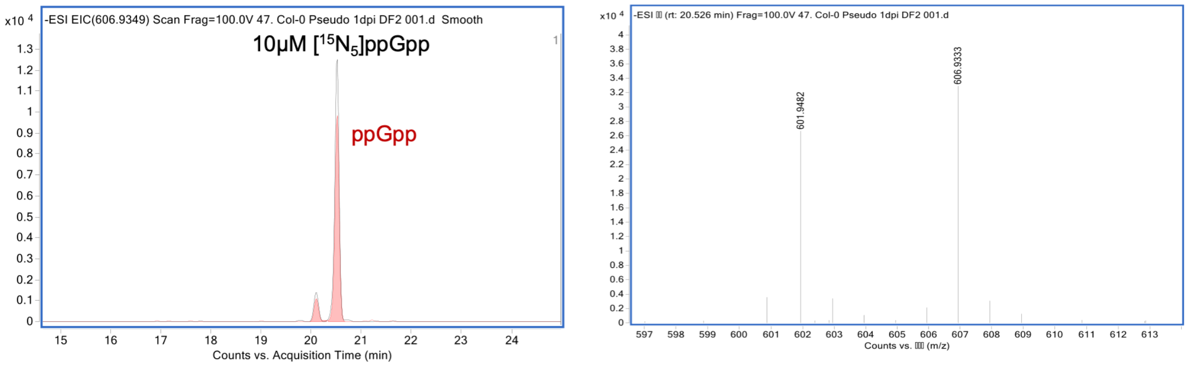


**Figure S4**: ppGpp analysis from Col-0 plant extracts infected with *PstDC3000* 1 dpi. Left: CE-MS electropherogram. Right: high-resolution mass spectrum of the analyte obtained on a Q-TOF System (6520, Agilent).

##### 3.2 Limit of Detection, Limit of Quantitation, Linearity

LOD and LOQ were determined via spiking of extracted plant material (150 mg FW) with heavy $^{15}N$ ppGpp standard (three biological repeats) on the QQQ System. LOD was defined as a signal to noise ratio (S/N) of above 3:1 and LOQ as S/N higher than 10:1. LOD was determined as 10 nM and LOQ was determined as 30 nM. One representative chromatogram of LOD and

LOQ is shown in figure S5 (LOD) and figure S6 (LOQ), respectively. Automatic noise selection (see figure S5 and S6, red) was artificial, so noise was selected manually (see figure S5 and S6, blue) and calculated by data analysis software.

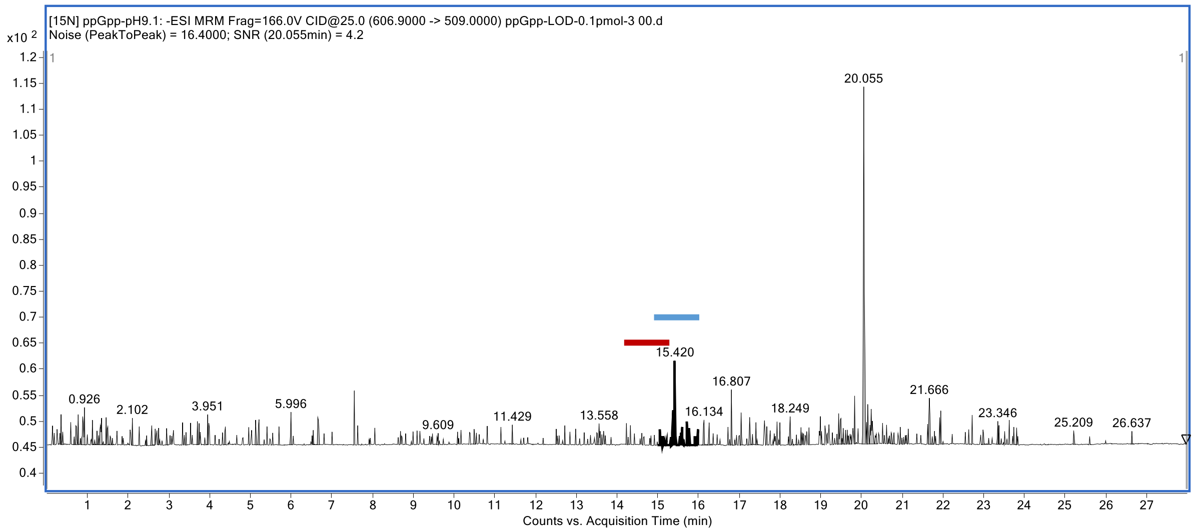


**Figure S5**: Determination of LOD

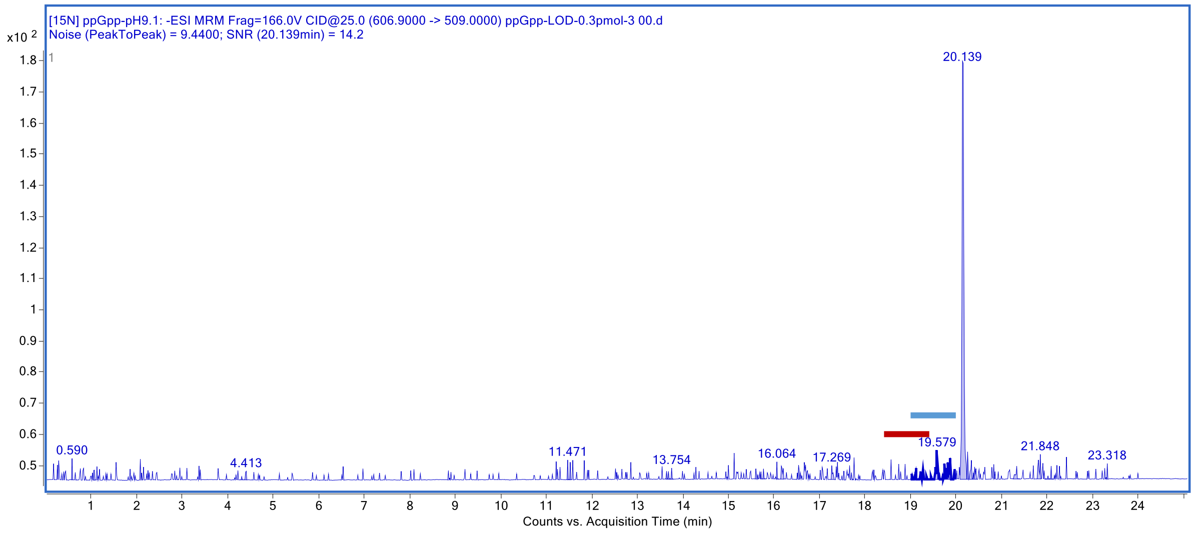


**Figure S6**: Determination of LOQ

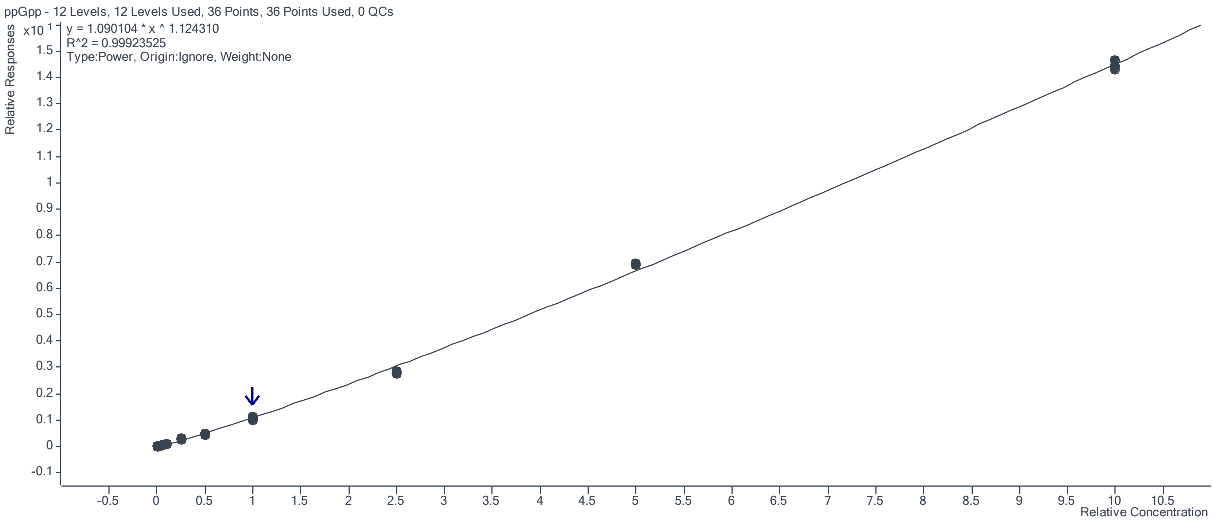


**Figure S7**: CE-MS analysis of a ppGpp dilution series from 0.1 µM- 400 µM. 12 levels, 36 points, $R^2$= 0.9992

#### 4. Bacterial and plant growth conditions

Arabidopsis seeds were surface sterilized and sown onto half-strength Murashige and Skoog (½ MS) medium supplemented with 1% sucrose. Plants were grown under 16 h/8 h day/night conditions at 22 °C/20 °C for 14 days. Light was provided by white LEDs ("True daylight", Polyklima). For *PstDC3000* treatment, plants were grown horizontally on plates. To conduct the flagellin assay, plants were grown vertically.*PstDC3000* were grown in King´s B medium + agarose for *PstDC3000* infection assay or in liquid King´s B with shaking at 220 rpm at 28°C for ppGpp extraction.

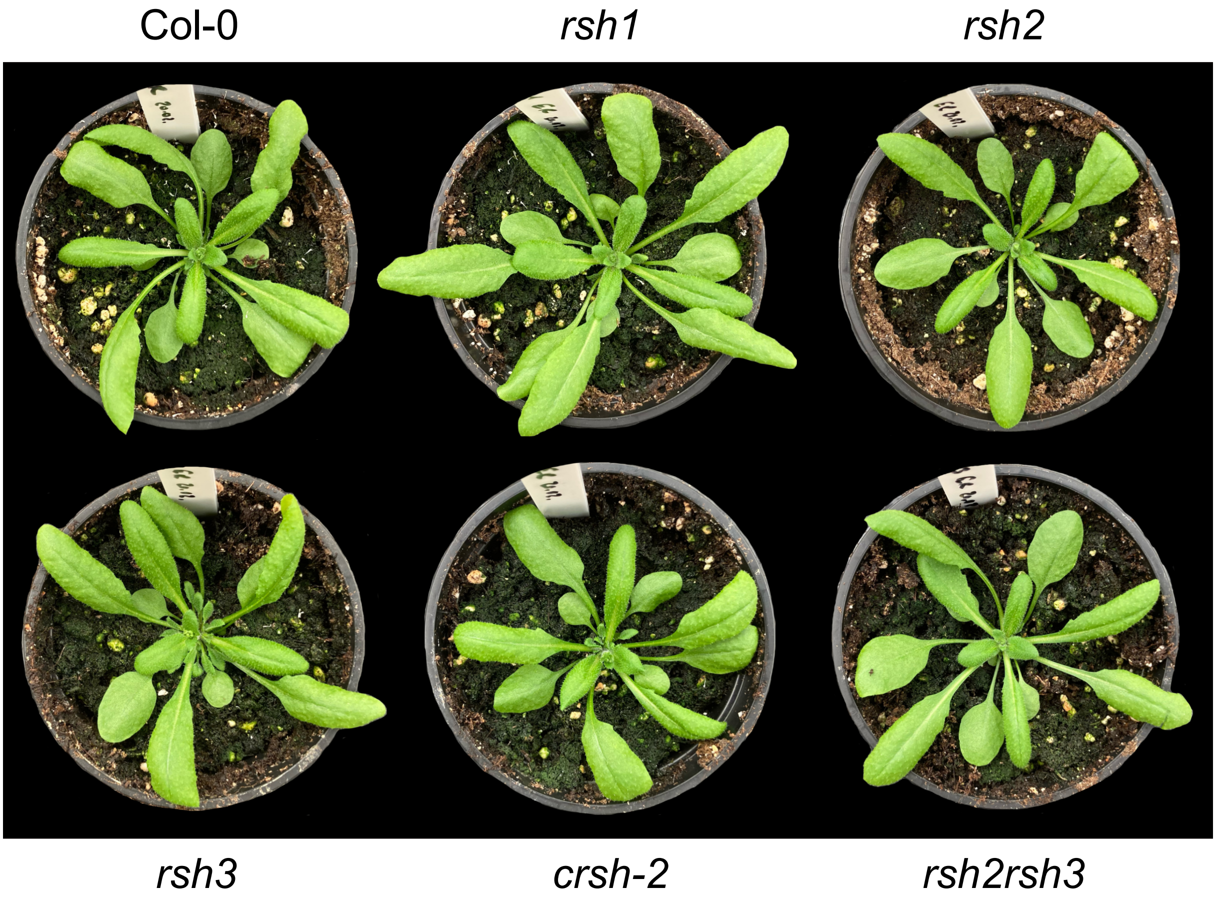


**Figure S8**: There is no obvious phenotype of mutant plants in comparison to wildtype plants (Col-0) under normal conditions.

#### **5. *PstDC3000* flood assay**

Infection of 14 days old *A. thaliana* seedlings was performed as described in Ishiga et al. (2017). *PstDC3000* was incubated at 28°C for 24-48h on plate. A single colony was incubated on fresh media and incubated at 28°C for 24-48h. Colonies were collected and resuspended in dd$H_2O$ to reach a final OD 0.1. Silwet L-77 was added to the suspension to a final concentration of 0.02%. $H_2O$ supplemented with 0.02% Silwet L-77 served as control. The suspension was dispensed onto ½ MS plates with seedlings and incubated for 3 min at room temperature. After incubation, the suspension was discarded and plates were sealed after drying. The seedlings were harvested 24h post infection (1 dpi) and surface sterilized in 5% $H_2O_2$. Samples were flash frozen in liquid nitrogen for ppGpp extraction.

#### 6. Gene expression analysis

Total RNA extraction of *PstDC3000* and flg22 treated plants was performed using the NucleoSpin RNA Plant and Fungi (Macherey-Nagel). For cDNA synthesis, 1 µg of RNA was treated with DNase I. The reverse transcription was performed according to the manufacturer´s instruction (Thermo Scientific™; RevertAid Reverse Transcriptase). SYBR Green reaction mix (Bio-Rad; SsoAdvanced Universal SYBR Green Supermix) was used in a Bio-Rad CFX384 real-time system for qPCR. *B-TUBULIN* was used as a reference gene.

#### 7. Chemical Synthesis

##### a) Synthetic procedures for [$^{15}$N]$_5$ – MSN

The [$^{15}$N]$_5$ – MSN analogues were synthesized in analogy to the procedures designed by Jessen et al. in 2019 (*Chem Commun* **2019,** *55* (37), 5339-5342) including minor modifications often related to the smaller reaction scale.

##### Synthesis of [$^{15}$N]$_5$ – pGp

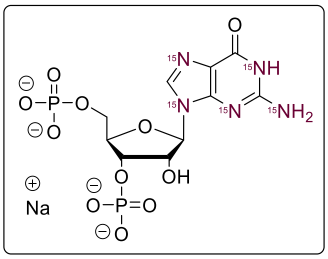

[$^{15}$N]$_5$ – guanosine dihydrate (23.0 mg, 79.8 µmol) was coevaporated using dry acetonitrile (3 x 2.0 ml). Afterwards, pyrophosphorylchloride (220 µl, 401 mg, 1.59 mmol, 20 eq.) was added at -35°C and the reaction mixture was stirred for 3 h at 0 °C. Subsequently, excess pyrophosphorylchloride was washed away with cold $Et_2O$ (3 x 10 ml) and the resulting crude product was hydrolyzed by adding $NaHCO_3$ – buffer (-2 °C, 1 M, 5.0 ml). The resulting solution was diluted with $H_2O$ (40 ml) and RNase $T_2$ (1.0 ku) was added. The mixture was incubated at 37 °C for 12 h. The crude product was purified by SAX ($NaClO_4$ – buffer). The product containing fractions were precipitated with a fourfold volume of $NaClO_4$ – solution (-20 °C, 0.5 M in acetone). The precipitate was washed with acetone (2 x 20 ml) and dried under high vacuum. The product (23.6 mg, 44.0 µmol, 55%) was isolated as Na-salt and white solid.

The cation was changed to TBA by using Dowex$H^+$.

**$^1$H-NMR** (400 MHz, $D_2O$, δ/ppm): 8.24 (dd, *J* = 11.0, 8.0 Hz, 1H), 5.97 (d, *J* = 7.1 Hz, 1H), 4.88 – 4.82 (m, 1H), 4.75 – 4.68 (m, 1H), 4.49 (br. s, 1H), 3.99 (dd, *J* = 4.0, 4.0 Hz, 2H). **$^{31}$P{$^1$H}-NMR** (162 MHz, $D_2O$, δ/ppm): 4.08, 3.82. **HRMS** (ESI) m/z for $C_{10}H_{13}O_{11}{}^{15}N_5P_2$ $[M-H_2]^{2-}$ calcd 222.9975 found 222.9974.

**Synthesis of [$^{15}$N]$_5$ – ppGpp**

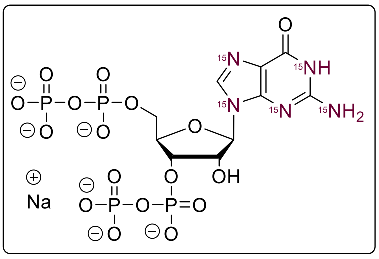


[$^{15}$N]$_5$ - pGp x 1.7 TBA (28.0 mg, 32.8 µmol) was dissolved in dry DMF (2.0 ml). ETT (21.2 mg, 164 µmol, 5.0 eq.) was added. Subsequently, a solution of $(FmO)_2P$-N*i*$Pr_2$ (90 %, 56.9 mg, 98.2 µmol, 3.0 eq.) in DMF (1.0 ml) was added and the resulting solution was stirred for 15 min at rt. The solution was cooled to -20 °C and *m*CPBA (77 %; 21.9 mg, 98.2 µmol, 3.0 eq.) was added. The solution was stirred for 10 min at -20 °C and DBU (300 µl) was added. The solution was stirred for 30 min at rt and the crude product was precipitated by the addition of $Et_2O$ (10 ml). The precipitate was separated by centrifugation and washed with $Et_2O$ (2 x 5 ml). The crude product was purified by SAX ($NaClO_4$ – buffer). The product containing fractions were precipitated with a fourfold volume of $NaClO_4$ – solution (-20 °C, 0.5 M in acetone). The precipitate was washed with acetone (2 x 20 ml) and dried over high vacuum. The product (16.8 mg, 22.7 µmol, 69%) was isolated as white solid.

**$^1$H-NMR** (400 MHz, $D_2O$, δ/ppm): 8.15 (dd, *J* = 11.1, 8.0 Hz, 1H), 6.00 (d, *J* = 6.4 Hz, 1H), 5.01 – 4.94 (m, 1H), 4.91 – 4.85 (m, 1H), 4.55 (br.s , *J* = 3.8 Hz, 1H), 4.29 – 4.16 (m, 2H). **$^{31}$P{$^1$H}-NMR** (162 MHz, $D_2O$, δ/ppm): -6.30 (d, *J* = 22.5 Hz), -7.73 (br.s), -10.87 (d, *J* = 15.6 Hz), -11.01 (d, *J* = 16.7 Hz). **HRMS** (ESI) m/z for $C_{10}H_{15}O_{17}{}^{15}N_5P_4$ $[M-H_2+Na]^-$ calcd 628.9168 found 628.9169.

**Synthesis of [$^{15}$N]$_5$ – pppGpp**

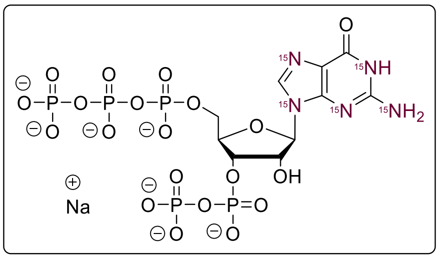


***Step 1:** synthesis of [$^{15}$N]$_5$ – ppGp*

[$^{15}$N]$_5$ - ppGpp (8.50 mg, 11.5 µmol) was dissolved in H2O (1.5 ml) and RNase T2 (1 ku) was added. The solution was acidified with HCl to pH 5.5 and incubated for 12 h at 37 °C. The crude product was purified by SAX ($NaClO_4$ – buffer). The product containing fractions were precipitated with a fourfold volume of $NaClO_4$ – solution (-20 °C, 0.5 M in acetone). The precipitate was washed with acetone (2 x 20 ml) and dried under high vacuum. The intermediate product [$^{15}$N]$_5$ – ppGp (4.55 mg, 7.13 µmol, 62 %) was isolated as Na-salt and white solid. The cations were exchanged to TBA by DowexH$^+$.

***Step 2:** synthesis of [$^{15}$N]$_5$ – pppGpp*

[$^{15}$N]$_5$ – ppGp x 2.6 TBA (7.80 mg, 6.76 µmol) was dissolved in dry DMF (1.0 ml). ETT (4.39 mg, 33.8 µmol, 5.0 eq.) was added. Subsequently, a solution of $(FmO)_2$P-N*i*Pr$_2$ (90 %, 11.7 mg, 20.3 µmol, 3.0 eq.) in DMF (1.0 ml) was added and the resulting solution was stirred for 15 min at rt. The solution was cooled to -20 °C and *m*CPBA (77 %; 4.53 mg, 20.3 µmol, 3.0 eq.) was added. The solution was stirred for 10 min at -20 °C and DBU (200 µl) was added. The solution was stirred for 30 min at rt and the crude product was precipitated by the addition of $Et_2O$ (10 ml). The precipitate was separated by centrifugation and washed with $Et_2O$ (2 x 5 ml). The crude product was purified by SAX ($NaClO_4$ – buffer). The product containing fractions were precipitated with a fourfold volume of $NaClO_4$ – solution (-20 °C, 0.5 M in acetone). The precipitate was washed with acetone (2 x 20 ml) and dried under high vacuum. The product (2.84 mg, 3.37 µmol, 50 %) was isolated as white solid.

**$^1$H-NMR** (400 MHz, $D_2O$, δ/ppm): 8.13 (dd, *J* = 11.0, 7.9 Hz, 1H), 6.00 (d, *J* = 6.4 Hz, 1H), 5.01 – 4.92 (m, 1H), 4.88 (dd, *J* = 5.8, 5.8 Hz, 1H), 4.54 (br. s, *J* = 3.9 Hz, 1H), 4.35 – 4.18 (m, 2H). **$^{31}$P{$^1$H}-NMR** (162 MHz, $D_2O$, δ/ppm): -5.74 (d, *J* = 22.3 Hz, 2P), -10.59 – -11.15 (m, 2P), -21.61 (dd, *J* = 19.8 Hz). **HRMS** (ESI) m/z for $C_{10}H_{16}O_{20}{}^{15}N_5P_5$ $[M-H_2]^{2-}$ calcd 342.9470 found 342.9473.

### 8. HRMS - spectra

#### *HRMS (ESI) analysis of [$^{15}N]_5$ - pGp*

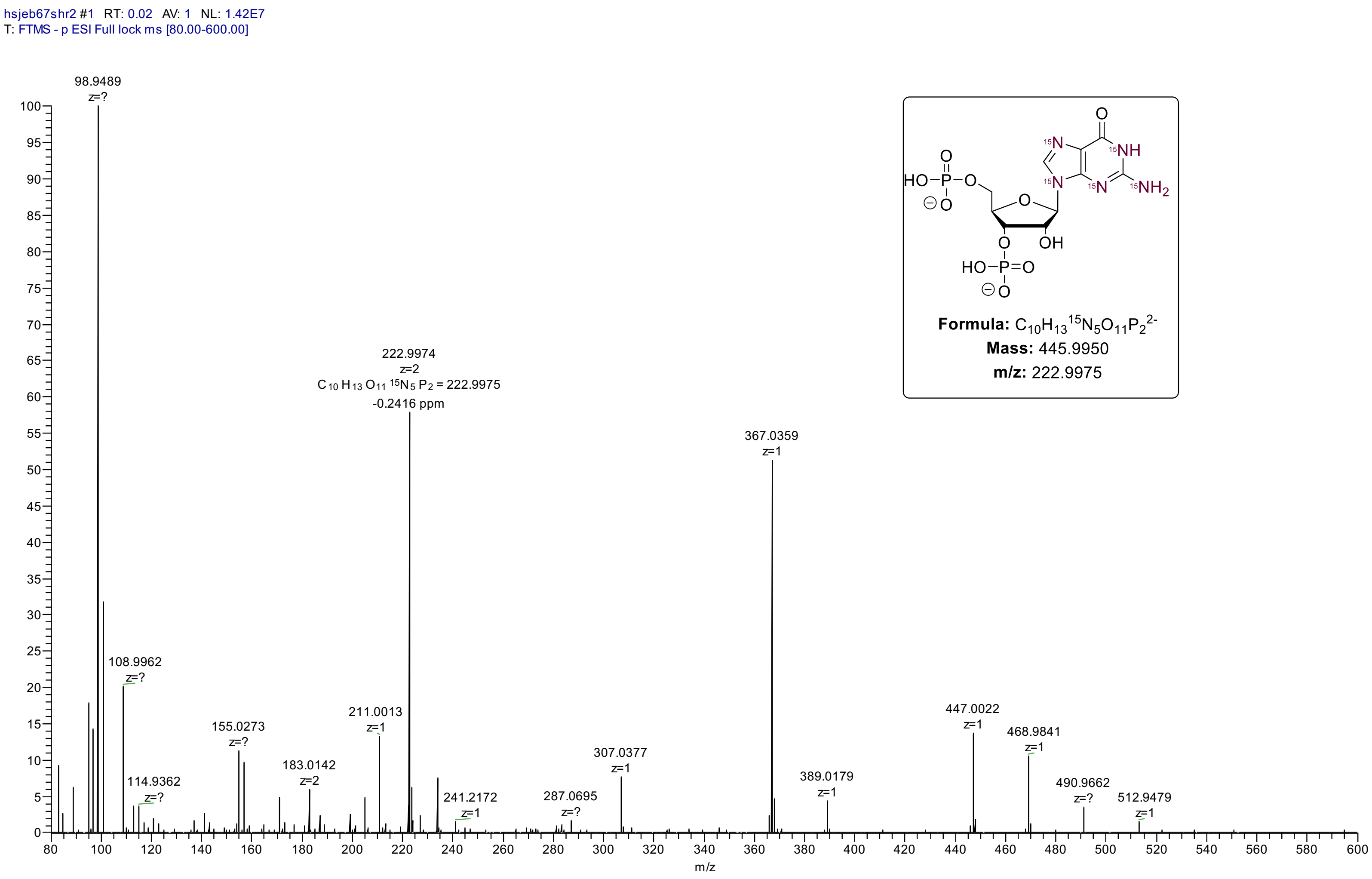

### HRMS (ESI) analysis of [$^{15}$N]$_5$ - ppGpp

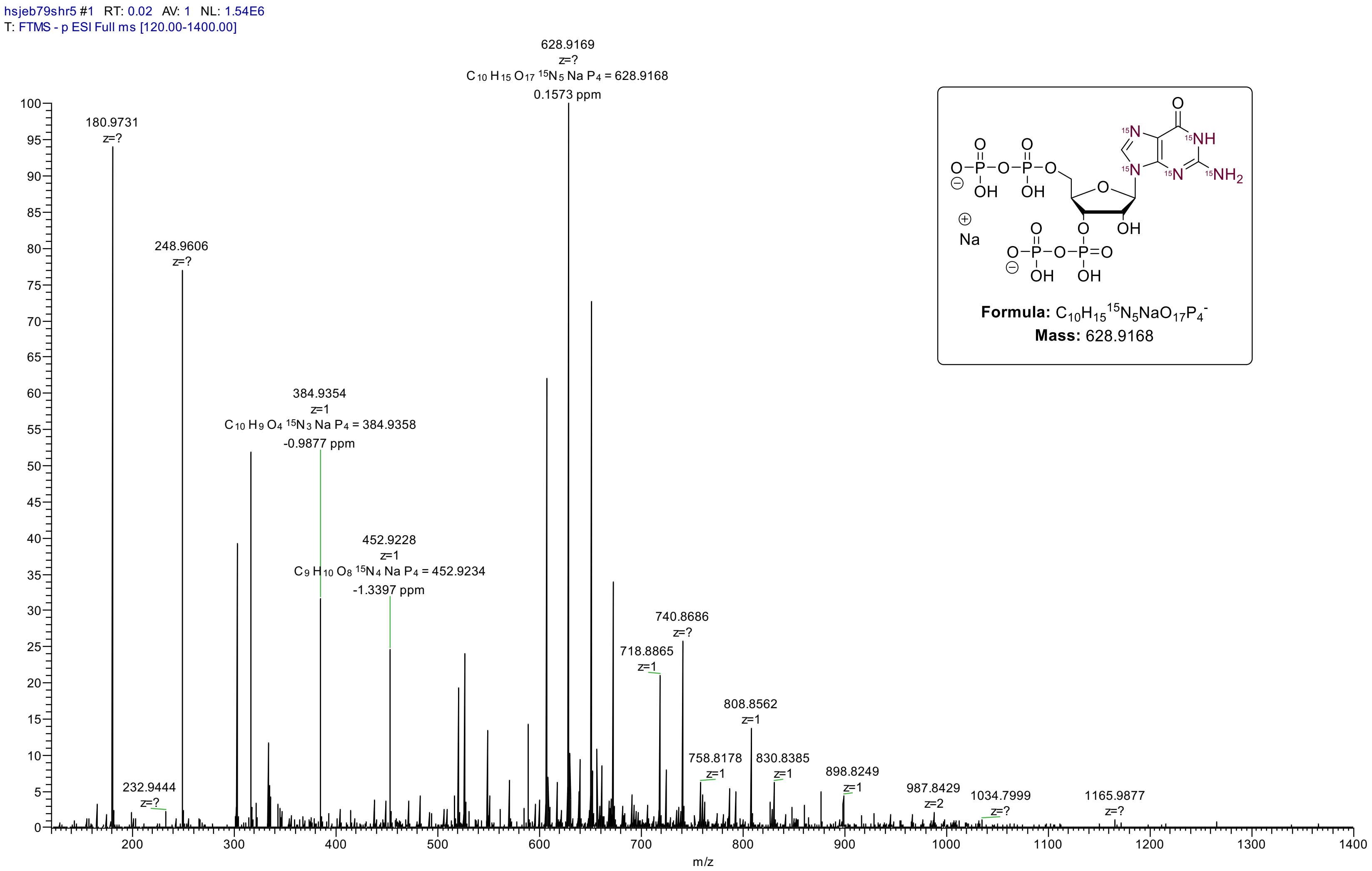

#### *HRMS (ESI) analysis of [$^{15}$N]$_5$ – pppGpp*

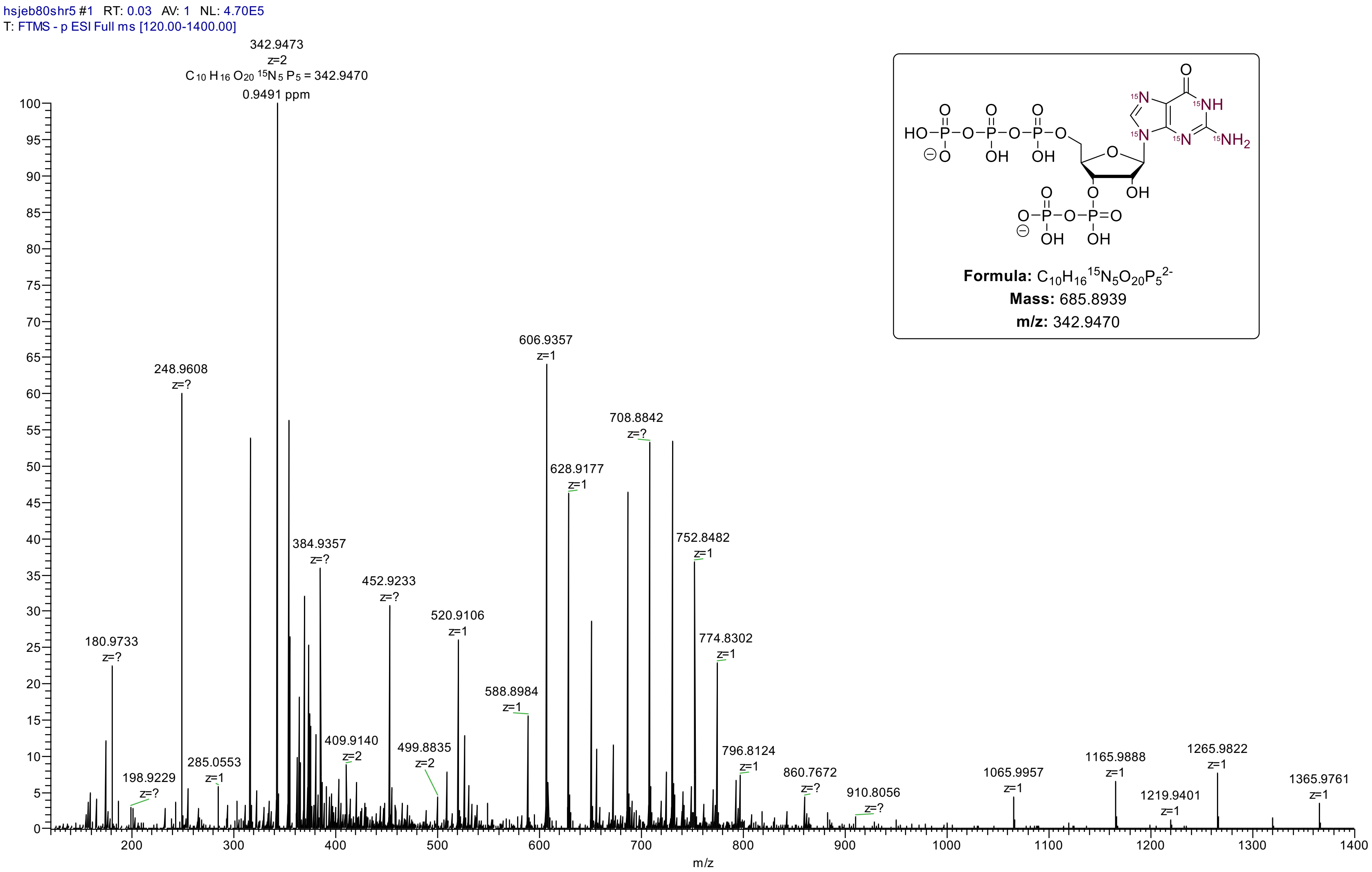

### 9. NMR - spectra

**Compound xx ($[^{15}N]_5$ - pGp), $^1H$ - NMR ($D_2O$, 400 MHz)**

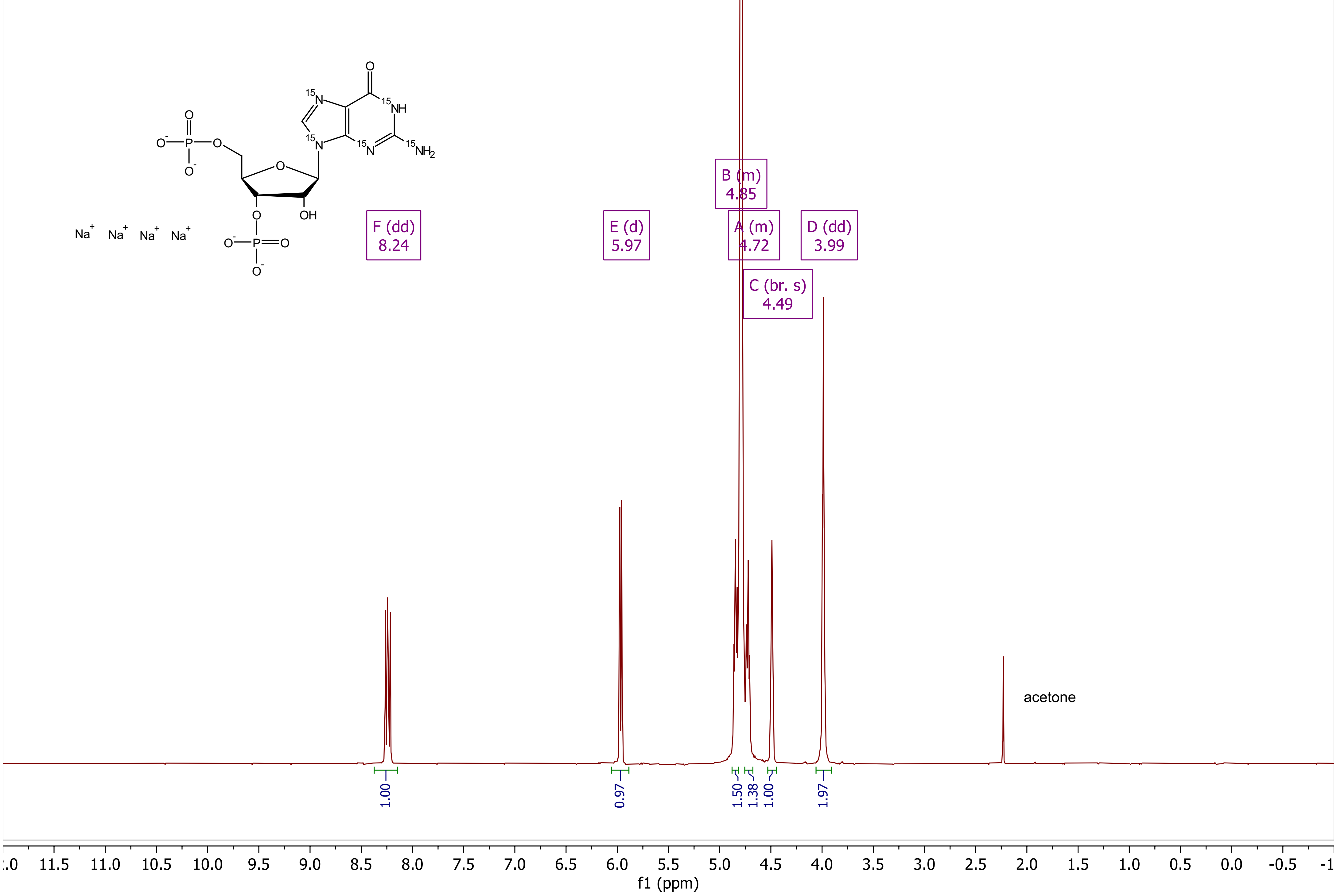

#### Compound xx ([$^{15}N$]$_5$ - pGp), $^{31}P$ {$^{1}H$} - NMR ($D_2O$, 162 MHz)

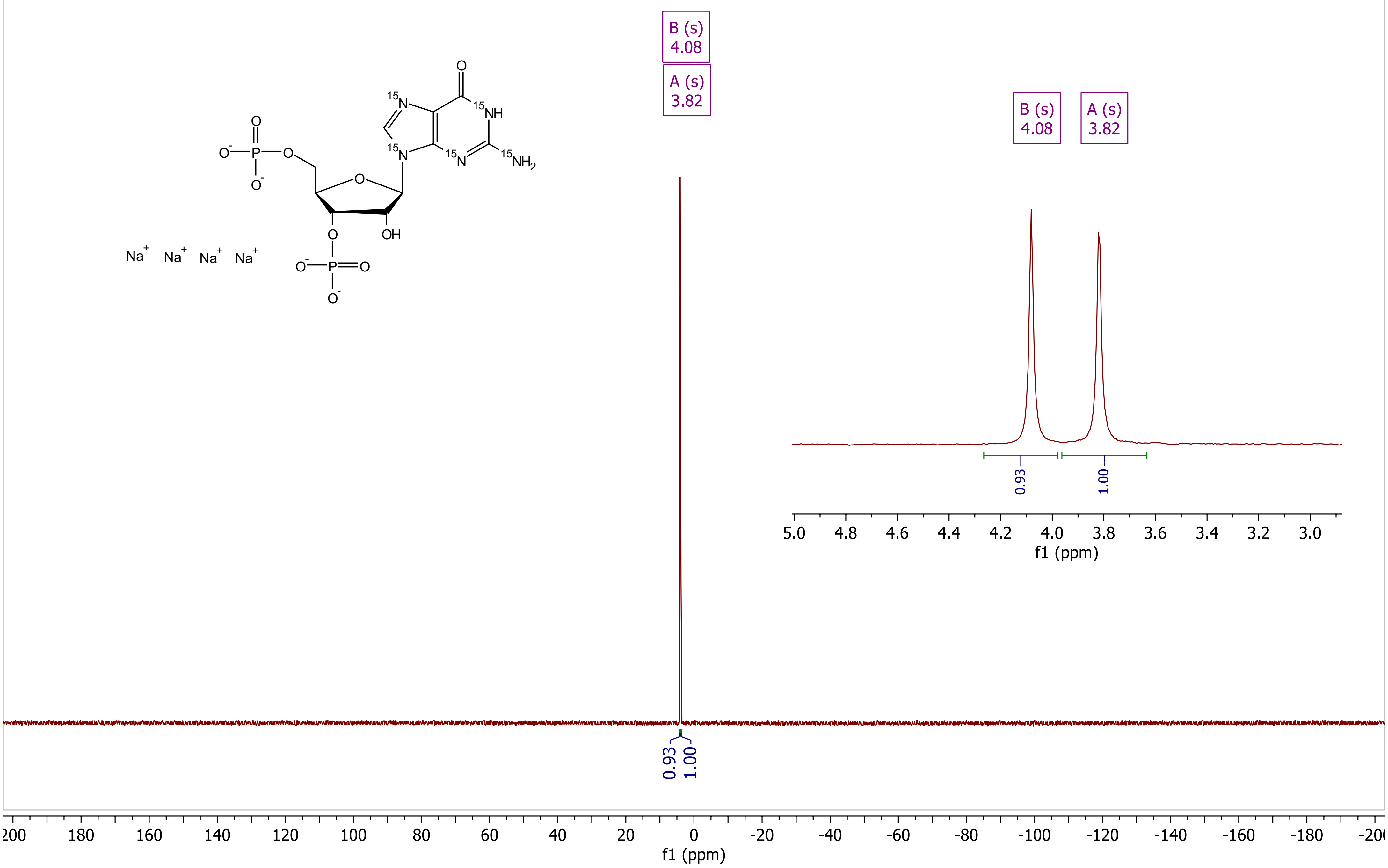

Compound xx ([$^{15}$N]$_5$ - ppGpp), $^1$H - NMR ($D_2O$, 400 MHz)

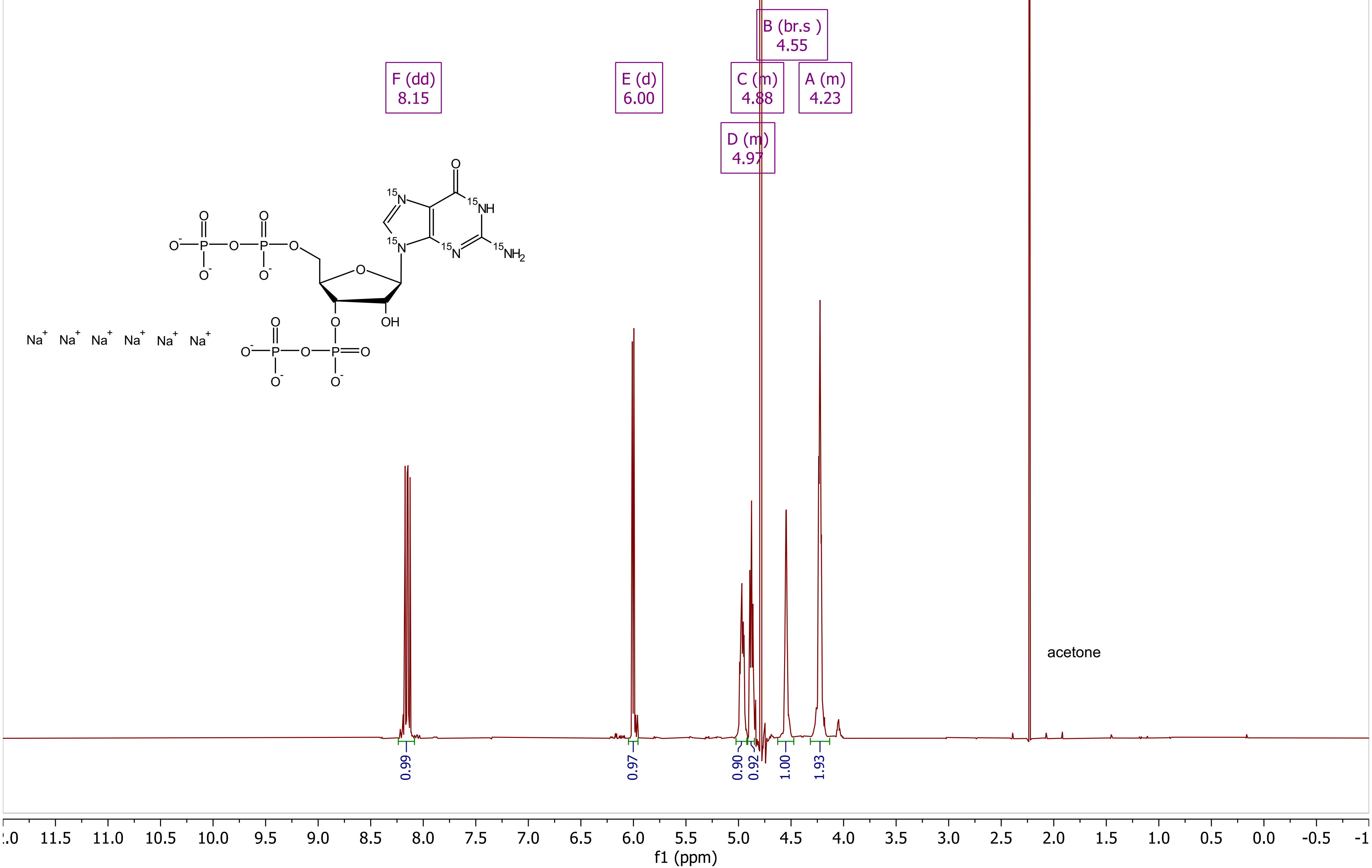

***Compound xx ([$^{15}$N]$_5$ - ppGpp), $^{31}$P {$^{1}$H} - NMR (D$_2$O, 162 MHz)***

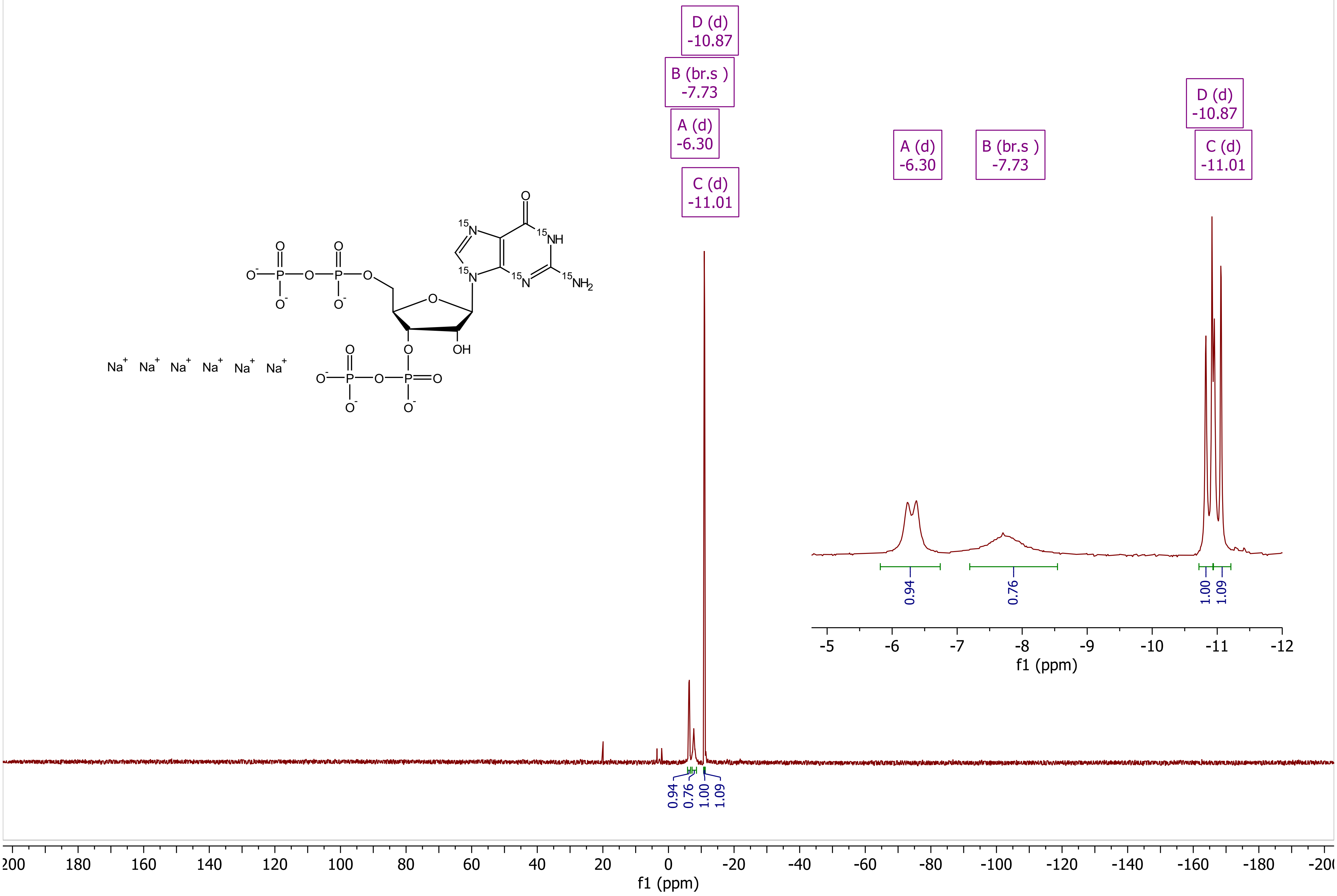

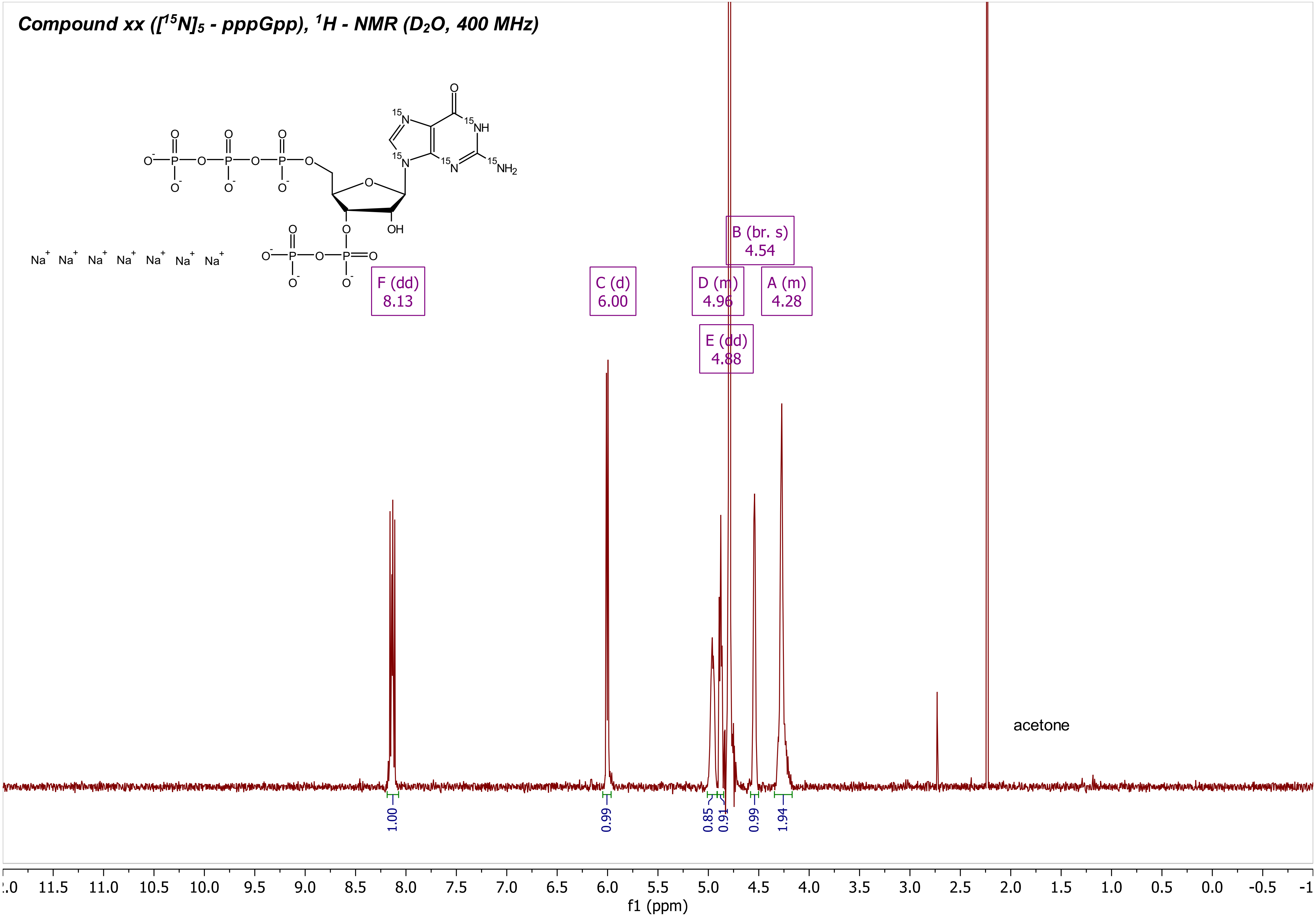
Compound xx ([15N]5 - pppGpp), 1H - NMR (D2O, 400 MHz)
Na+ Na+ Na+ Na+ Na+ Na+ Na+
15N
15NH
15N
15N
15NH2
O
OH
F (dd)
8.13
C (d)
6.00
D (m)
4.96
E (dd)
4.88
B (br. s)
4.54
A (m)
4.28
acetone
1.00
0.99
0.85
0.91
0.99
1.94
f1 (ppm)

**Compound xx ([$^{15}$N]$_5$ - pppGpp), $^{31}$P {$^1$H} - NMR (D$_2$O, 162 MHz)**

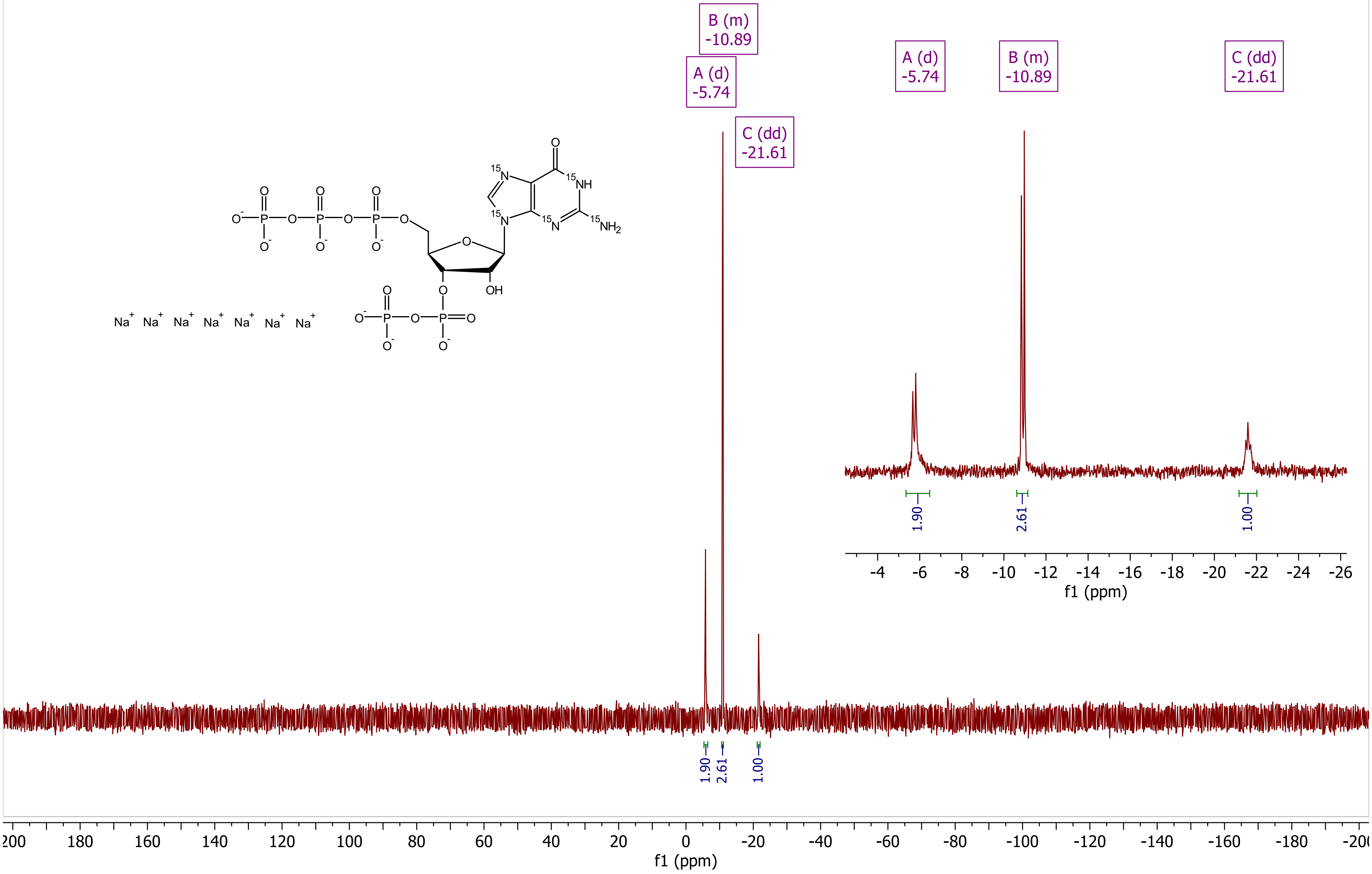
